# Codon-specific ribosome stalling reshapes translational dynamics during branched-chain amino acid starvation

**DOI:** 10.1101/2025.03.27.645654

**Authors:** Lina Worpenberg, Cédric Gobet, Felix Naef

## Abstract

Cells regulate protein synthesis in response to fluctuating nutrient availability through coordinated mechanisms that affect both translation initiation and elongation. Branched-chain amino acids (BCAAs)—leucine, isoleucine, and valine—are essential nutrients with a significant impact on cellular physiology. However, how their simultaneous depletion affects translation remains largely unclear. Here, we examined the immediate effects of short-term BCAA limitation on translational dynamics in mammalian cells. We performed RNA sequencing (RNA-seq) and ribosome profiling (Ribo-seq) on NIH3T3 cells subjected to single, double, or triple BCAA deprivation. Our analyses revealed increased ribosome dwell times in all starvation conditions, with pronounced stalling at all valine codons during valine and triple starvation, while leucine and isoleucine starvation produced milder, codon-specific effects. Notably, we could show that isoleucine starvation-specific stalling largely diminished under triple starvation, likely due to early elongation bottlenecks at valine codons. Correlating these stalling events with tRNA charging levels revealed distinct tRNA isoacceptor regulation in each starvation condition. In addition, integrating quantitative proteomics showed that many proteins downregulated under BCAA deprivation harbor stalling sites, suggesting that compromised elongation contributes to decreased protein output. Together, these findings suggest that differential ribosome stalling under BCAA starvation reflects a balance between amino acid supply, tRNA charging dynamics, and stress-response signaling, illustrating that codon choice shapes the translational landscape under nutrient limitation.

## Introduction

Branched-chain amino acids (BCAAs) — leucine, isoleucine, and valine — are essential nutrients involved in diverse cellular and physiological processes, including protein synthesis, metabolism, cellular signaling, brain function, and immune responses ^1–5^. Because mammalian cells cannot synthesize these amino acids, they rely on external sources to maintain proteostasis and meet metabolic demands. Accordingly, fluctuations in BCAA levels can have strong implications for health and disease ^6–8^. Elevated BCAA concentrations are associated with obesity, insulin resistance, and type 2 diabetes mellitus ^9–12^, while low BCAA plasma levels are correlated with the prevalence of liver cirrhosis and hepatic encephalopathy^13^, and shifts in BCAA metabolism have been linked to tumor growth, progression, and survival ^14–17^.

Translation is one of the most energy and resource consuming processes within dividing cells, and thus, is tightly regulated in response to nutrient availability ^18^. Two key pathways govern this response: The mechanistic target of rapamycin complex 1 (mTORC1) pathway and the Integrated Stress Response (ISR). The modulation of both pathways aims to reduce global translation initiation to maintain cellular homeostasis and to help cells cope with stress. Under normal conditions, mTORC1 promotes translation initiation by phosphorylating substrates such as eukaryotic translation initiation factor 4E-binding protein 1 (4E-BP1) and ribosomal protein S6 kinase (S6K) ^19–21^, thereby promoting the translation of transcripts which contain a terminal oligopyrimidine tract (TOP) that encode ribosomal and other translational machinery components ^22,23^. When amino acid levels are limited, mTORC1 activity drops, causing reduced global translation initiation, downregulated translation of TOP transcripts, and partial reprogramming of gene expression. Meanwhile, the General Control Nonderepressible 2 (GCN2) kinase, a central component of the ISR, responds to increased levels of uncharged tRNAs that accumulate when particular amino acids are deficient ^24,25^. This triggers phosphorylation of eukaryotic translation initiation factor 2 subunit 1 (eIF2α), which suppresses cap-dependent initiation while selectively promoting translation of stress-response genes ^26–28^.

Although many studies have focused on how these pathways downregulate translation initiation, it is increasingly evident that amino acid availability also controls translation elongation. A steady supply of aminoacyl-tRNAs is required for continuous elongation, and a shortage of even one amino acid can trigger codon-specific ribosome stalling ^29^. A recent example shows that leucine starvation leads to reduced charging of tRNA^Leu^_UAA_, which causes ribosome stalling at UUA codons and promotes mRNA frameshifting and protein misfolding ^30^. This selective charging of tRNA isoacceptors adds another layer of regulation, whereby some synonymous codons become more prone to stalling than others ^31,32^. Besides the availability of charged tRNAs, the sequence context of mRNA itself plays a crucial role in modulating ribosome dynamics during translation as both codon composition and secondary mRNA structures influence translation elongation. Stable secondary structures, such as stem-loops or pseudoknots, can create physical barriers to ribosome progression leading to transient pausing or stalling ^33^. Likewise, clusters of rare codons can slow translation by limiting the availability of cognate tRNAs, further affecting elongation kinetics ^34^. These pauses not only influence translation efficiency but also impact co-translational protein folding and may serve as regulatory checkpoints, particularly under conditions of nutrient limitation. This phenomenon underscores the complexity of codon-level control under nutrient stress.

Recent technical advances, including ribosome profiling (Ribo-seq), have made it possible to study translation at single-codon resolution ^35^. By isolating and sequencing ribosome-protected mRNA fragments (RPFs), one can directly observe where ribosomes pause during normal growth or under stress. In particular, codon-specific ribosome “dwell times” (DT) can be inferred from Ribo-seq data, serving as an indicator of how long the ribosome remains at each codon ^36^.

However, while previous research has highlighted the impact of individual amino acid shortages on the translation dynamic and that the effects vary depending on the specific amino acid being limited ^37–39^, a comprehensive understanding of how simultaneous BCAA limitation influences translation is lacking. Here, we address this gap in the field by systematically employing Ribo-seq upon single and simultaneous starvation of BCAAs to assess global translational output and codon-specific dwell times. This approach allows us to identify unique stalling patterns associated with each BCAA deprivation. Complementary quantitative proteomics experiments allowed us to correlate these translational modifications with changes in protein abundance. Additionally, tRNA charging assays elucidated the mechanistic relationship between amino acid depletion, uncharged tRNA isoacceptors, and ribosomal pausing. Through the integration of these multi-omics datasets, we demonstrated that BCAA starvation induces diverse translational control patterns, reflecting the interplay of mTORC1 and GCN2 pathways, tRNA isoacceptor availability, and transcript codon usage. Additionally, our analysis highlights a positional effect, indicating that translational regulation is not only influenced by the codon composition of transcripts but also by the location of codons within the mRNA sequence. This study uncovered a previously underappreciated layer of complexity in BCAA-specific translational regulation, setting the stage for future investigations into the precise mechanisms that emerge when valine, isoleucine, and leucine are concurrently limited.

## Material and Methods

### Cell culture and amino acid starvation

NIH3T3 mouse fibroblast cells were cultured in DMEM (high glucose, pyruvate; Gibco) supplemented with 10% FBS (qualified, USDA-approved regions; Gibco), and 100 U/ml penicillin-streptomycin (Gibco). Cells were maintained at 37°C in a humidified incubator with 5% CO_2_. Amino acid starvation experiments were conducted using custom DMEM/F-12 media (Gibco; SKU11320 modified) supplemented with 10% dialyzed FBS (Gibco) and lacking specific BCAAs — leucine, isoleucine, and valine — as well as asparagine and aspartate. L-Asparagine monohydrate (7.5 mg/l) and L-aspartic acid (6.65 mg/l) were added back to all media. Leucine (59.05 mg/l), isoleucine (54.47 mg/l), and valine (52.85 mg/l) were supplemented in specific combinations to create the following media: Complete medium (containing all amino acids at standard concentrations), Leucine starvation (medium lacking leucine), Isoleucine starvation (medium lacking isoleucine), Valine starvation (medium lacking valine), Double starvation (medium lacking both leucine and isoleucine), Triple starvation (medium lacking leucine, isoleucine, and valine). The amino acids were purchased in powdered form from Sigma-Aldrich. Cells were washed once with 1xPBS (Gibco) before switching to the respective starvation medium for 6 hours.

### Ribo-seq & RNA-seq

Cells were washed with cold PBS (Gibco) supplemented with 100 µg/ml cycloheximide (Sigma-Aldrich), lysed in 200 µL polysome lysis buffer (20 mM Tris-HCl pH 7.4, 150 mM NaCl, 5 mM MgCl2, 1% Triton X-100, 5 mM DTT, 100 µg/ml CHX, 25 U/mL TURBO DNase (Invitrogen), 10 U/µL murine RNase inhibitor (NEB), and protease/phosphatase inhibitors (Sigma-Aldrich). Cells were scraped down, and transferred into microcentrifuge tubes. After incubation on ice for 10 minutes, lysates were passed through a 26G needle 10 times and centrifuged at 20,000g for 10 minutes at 4°C. Supernatants were collected and quantified by NanoDrop. Ribosome-protected fragments (RPFs) were obtained by adding 43 U/OD260 RNase I (Invitrogen) at 24°C for 45 minutes. The reaction was stopped with 5 µL SUPERase In RNase inhibitor (Invitrogen) and placed on ice. Digested samples were purified using Microspin S-400 HR columns (Cytiva) equilibrated with polysome buffer (20 mM Tris-HCl pH 7.4, 150 mM NaCl, 5 mM MgCl₂, 1 mM DTT, and 100 µg/ml cycloheximide) for 2 minutes at 2400 rpm, followed by phenol-chloroform extraction and ethanol precipitation. Precipitated RNA was resuspended in water and quantified. For size selection, 15% Novex TBE-Urea polyacrylamide gels (Invitrogen) were pre-run for 20 minutes at 200V. Samples were mixed with Novex TBE-Urea sample buffer (Invitrogen), heated at 90°C for 2 minutes, and loaded onto the gels, alongside RNA size markers (NEB). Gels were run at 170V for approximately 70 minutes and stained with SYBR Gold gel stain (Invitrogen). Gels were excised under blue light and fragments between 25-36 nucleotides purified. To extract the RNA from gel fragments, a 0.5 mL microcentrifuge tube was pierced at the bottom with an 18G needle, the cap was removed, and the tube was placed inside a 1.5 ml microcentrifuge tube. The gel fragment was placed inside the prepared 0.5 ml tube and centrifuged for 2 minutes at full speed to collect the gel debris in the lower tube. 360 µl of water was added to the gel debris, and incubated at 70°C for 10 minutes. The resulting gel slurry was transferred into a microfuge tube spin filter (Corning Costar Spin-X) and centrifuged for 3 minutes at full speed. The filtrate was collected in a new tube, and 40 µL of 3 M sodium acetate, pH 4.5 was added. Subsequently, 1.5 µl of GlycoBlue (Invitrogen) was added, followed by 700 µL of isopropanol. The mixture was incubated overnight at −20°C for precipitation. RNA was collected by centrifugation, pellets were washed with 80% ethanol, dried, and resuspended in water for further processing. rRNA was depleted by hybridizing RNA samples with complementary biotinylated oligonucleotides (see Supplementary Material and Methods) and hybridized rRNA was removed using MyOne Streptavidin C1 Dynabeads (Invitrogen). Samples were purified using a RNA Clean & Concentrator kit (Zymo). End repair was performed by incubating RNA with T4 PNK (NEB) at 37°C for 1 hour, followed by ATP addition and another hour of incubation. Clean-up was performed using the RNA Clean & Concentrator kit (Zymo). Libraries were prepared using the NEXTflex Small RNA-Seq Kit v3 (Revvity) according to the manufacturer’s protocol. RNA from the same experimental conditions was used for RNA-seq analysis. For this, RNA was extracted from the polysome lysates before RNase I digestion by phenol-chloroform extraction and ethanol precipitation. Total RNA was submitted to the EPFL Gene Expression Core Facility (GECF) for mRNA library preparation using NEBNext Ultra II Directional RNA Library Prep with PolyA selection. RNA-seq libraries were sequenced on a Novaseq6000 generating paired-end (50/50) reads and Ribo-seq libraries were sequenced on a Nextseq500 system generating single-end 75 reads.

### RNA-seq and Ribo-seq data processing

For RNA-seq, paired-end reads were aligned to the Ensembl mouse GRCm38 primary assembly reference genome using STAR (v2.7.11a)^40^. The corresponding Ensembl GTF annotation (GRCm38.100) was used to build the genome index, and gene-level counts were generated with the --quantMode GeneCounts option. Reads mapping in the sense orientation relative to annotated transcripts were retained. On average, approximately 51 million reads per sample were uniquely mapped and quantified.

For Ribo-seq, adapter sequences were trimmed from raw Ribo-seq reads using cutadapt^41^ with the parameter -m 10 and the adapter sequence TGGAATTCTCGGGTGCCAAGG. Using an in-house Perl script, we then removed duplicate reads carrying identical sequences and UMIs (four random nucleotides at both 5′ and 3′ ends). The processed reads were first aligned to a combined mouse and human rRNA/tRNA reference (from UCSC genome browser) using STAR to remove and quantify rRNA/tRNA contaminants, then mapped to the same GRCm38 reference genome used for RNA-seq. To select ribosome-protected fragments, only reads between 25 and 35 nucleotides in length with a unique alignment ([NH]==1) were retained using samtools. Feature distribution was assessed with RSeQC read_distribution.py ^42^ using the mm10_GENCODE_vm25.bed file on the filtered BAM files. Finally, gene-level counts were primarily generated with htseq-count using the intersection-strict mode ^43^ from the same filtered BAM file, as well as the same GTF annotation used for the RNA-seq analysis. For a comparative analysis, an additional count matrix was generated including multi-mapped reads (htseq-count --nonunique all -a 0). This multimapping dataset is used in Supplementary Figure 5 for comparison, while all other analyses are based on uniquely mapped reads.

Only protein-coding genes were retained for further analysis and genes without gene symbol annotation or with fewer than 10 reads were removed. Differential gene analysis was performed in R using edgeR: Separate DGEList objects were created for RNA-seq and Ribo-seq datasets and normalized, followed by estimation of dispersion parameters using estimateDisp. A design matrix was constructed and genewise quasi-likelihood negative binomial GLMs were fitted. Differential expression (DE) testing was performed using glmQLFTest for each contrast of interest. For each condition, DE results were extracted, and p-values were adjusted for multiple testing using the Benjamini–Hochberg procedure. Ribosome densities (RD) were calculated as the ratio of the RPM of Ribo-seq/RNA-seq.

Dwell time modeling. For ribosome dwell time inference and gene ribosome density profiles, the Ribo-seq de-duplicated reads were input to the Ribo-DT pipeline ^44^ using similar reference genomes as above. Parameters were set as follows: a lower bound read size (L1) of 26 nucleotides, an upper bound (L2) of 35 nucleotides, library strandness specified as “pos_neg”, and a filter threshold of 10 reads per gene. Single dwell time output table was used for all subsequent analysis. Intermediate “.count” Ribo-DT output files were used to generate ribosome density profiles. Mean DTs for each experimental condition were calculated by averaging triplicate values. Unless specified otherwise, codon-level DTs were filtered to include specific ribosomal positions (“-2”, “-1”, “E”, “P”, “A”, “3”, “4”) and summarized as mean values per codon. For statistical analysis, codon-specific ANOVA models were applied to compare DTs across conditions, followed by post-hoc Tukey′s Honest Significant Difference (Tukey HSD) tests to identify significant differences between experimental conditions and the control.

### Polysome profile generation and analysis

Polysome lysates were prepared as described above for Ribo-seq analysis. Sucrose gradients (5%–50% in polysome buffer: 20 mM Tris-HCl pH 7.4, 150 mM NaCl, 5 mM MgCl₂, 1 mM DTT, and 100 µg/ml cycloheximide) were prepared using a gradient maker (Biocomp). Lysates were loaded onto the gradients and centrifuged at 32,000 rpm for 2.5 hours at 4°C in an ultracentrifuge (Beckman Coulter). Polysome profiles were generated by fractionating the gradients with continuous UV monitoring at 260 nm using an automated gradient fractionation system (Biocomp). The raw data was further processed with R: Raw polysome profiles were smoothed using moving average filters to reduce noise. For alignment, manually selected reference points representing key structural features of the profiles (anchor points; monosome and polysome peaks, and local maxima and minima) were defined for each sample. To standardize baseline values, each profile was shifted such that the overall minimum intensity was set to zero. Following this, the defined anchor points were used to perform a linear time axis interpolation to correct for run-to-run variation in migration patterns. After alignment, the intensity of each profile was normalized to its total area under the curve (AUC), enabling cross-sample comparisons, and AUC for polysome and monosome area was calculated. Polysome-to-monosome ratios (P/M) were calculated and values were further normalized to the control condition. Statistical analyses were performed using unpaired t-tests. All results are presented as mean ± SEM, with statistical significance indicated.

### Intracellular amino acid measurement

Cells were incubated in the respective starvation media for 6 hours, as detailed above. After treatment, cells were washed twice with ice-cold phosphate-buffered saline (PBS) to remove extracellular amino acids. Plates were then flash-frozen on dry ice to preserve intracellular metabolites. Frozen samples were submitted to the Metabolomics Unit at the University of Lausanne (UNIL) for analysis. Amino acids were quantified using a stable isotope dilution LC-MS approach, following established protocols ^45^. Each condition was analyzed in five biological replicates.

### Western blot analysis

Protein lysates were prepared by resuspending cell pellets in a hot SDS lysis buffer (2% SDS, 50 mM Tris-HCl, pH 7.5, 1 mM EDTA) and heating samples at 95°C for 5 minutes. Afterwards, samples were centrifuged for 5 minutes at max speed and supernatant transferred to a new tube. Protein concentrations were measured using a NanoDrop spectrophotometer (Thermo Fisher Scientific). Protein lysates were mixed with NuPAGE LDS sample buffer (Invitrogen) and 50 mM DTT. Equal amounts of protein (8 µg per lane) were separated on 8%-12% gradient SDS-PAGE gels (SurePAGE, Bis-Tris; GeneScript) and transferred onto PVDF membranes using the iBlot 2 Dry Blotting System with PVDF transfer stacks (Invitrogen). Membranes were blocked in 5% BSA in PBS-T (PBS with 0.1% Tween-20) for 1 hour at room temperature. Primary antibody incubation was performed overnight at 4°C in 5% BSA in PBS-T. After washing three times with PBS-T, membranes were incubated with HRP-conjugated secondary antibodies for 1 hour at room temperature. Chemiluminescent signals were detected using the Fusion FX imaging system (Vilber). Protein band intensities were quantified using ImageJ software (NIH), and target protein levels were normalized to the corresponding loading control. Results represent the mean ± standard deviation from three independent biological replicates. Significance was determined by unpaired t-tests.

### RT-qPCR analysis

Total RNA was extracted using TRIzol reagent (Thermo Fisher Scientific) according to the manufacturer′s instructions. RNA concentration and purity were assessed using a NanoDrop spectrophotometer (Thermo Fisher Scientific). Complementary DNA (cDNA) synthesis was performed with 1 µg of RNA using M-MLV Reverse Transcriptase (Promega) and random hexamer primers in a 20 µL reaction. qPCR was carried out using a SYBR Green-based qPCR master mix (Promega) according to the manufacturer′s protocol on a QuantStudio 7 Real-Time PCR System (Applied Biosystems). Cycling conditions included an initial denaturation at 95°C for 2 minutes, followed by 40 cycles of denaturation at 95°C for 15 seconds and annealing/extension at 60°C for 1 minute. Relative gene expression levels were determined using the ΔΔCt method, normalizing target gene expression to Gapdh as housekeeping gene. Data represent the mean ± standard deviation from biological replicates, with each sample analyzed in technical duplicates. Used primers can be found in the Supplementary Material and Methods.

### OPP incorporation assay

OPP (O-propargyl-puromycin) incorporation was used to assess protein synthesis following 6 hours of different starvation conditions. Approximately 8,000 cells per well were seeded into black 96-well plates with glass bottoms (Greiner Bio-One) the day before the assay. OPP was added directly to the culture medium at a final concentration of 25 µM (prepared from a 2.5 mM stock solution in DMSO, diluted 1:100) and incubated with cells for 20 minutes. After incubation, the medium was removed, and cells were washed twice with PBS. Cells were fixed with 4% PFA in PBS (ABCR) for 5 minutes at room temperature. Following fixation, cells were permeabilized with 0.1% Triton X-100 in PBS for 5 minutes and subsequently washed twice with PBS and once with PBS containing 3% BSA. Click chemistry was performed by incubating cells with 150 µl of reaction mix containing 1× PBS, 500 µM CuSO₄, 2 µM 5-TAMRA-Azide (Jena Bioscience), and 50 mM L-ascorbic acid (Sigma Aldrich) for 45 minutes at room temperature. After incubation, cells were washed sequentially: once with 0.5 mM EDTA in PBS, twice with PBS containing 3% BSA, and finally with PBS containing Hoechst 33342 (1:10000 dilution, Thermo Scientific) for nuclear staining. Fluorescence images were acquired using the Operetta High-Content Imaging System (PerkinElmer). OPP incorporation and nuclear staining were quantified for individual cells using the Harmony software (PerkinElmer). Local background fluorescence was subtracted and, to account for autofluorescence, data from unstained samples were subtracted from all experimental conditions. Outliers were identified and removed using the interquartile range (IQR) method, excluding values outside 1.5 × IQR. Fluorescence intensities were further normalized to the mean of control samples. Pairwise comparisons to control samples were performed using Bonferroni-adjusted unpaired t-tests.

### TMT-Based quantitative proteomics

Cells were seeded in 6-well plates one day prior to treatment. The medium was replaced with 2 ml of deprivation medium, and cells were incubated for 6 hours. After incubation, cells were trypsinized, centrifuged, and washed once with PBS. The cell pellets were resuspended in 100 µL of lysis buffer containing 2% SDS, 50 mM Tris (pH 8.0), protease, and phosphatase inhibitors. Samples were boiled for 5 minutes and passed through a 26G needle 10 times for homogenization, followed by centrifugation at maximum speed for 5 minutes to remove debris. Protein concentrations were determined by BCA assay. Equal amounts of protein from each sample were processed for TMT-based quantitative proteomic analysis at the EPFL Proteomics Core Facility. Samples were digested by FASP (Filter Aided Sample Preparation) following the standard procedure with minor modifications ^46^. Protein samples containing 20 µg were prepared in 2% SDS, 100mM Tris-HCl pH 8.0 and deposited on conditioned filter devices (30K). Devices were centrifuged at 10’000 x g for 30 min. until complete dryness and then washed twice with 200 µL of Urea solution (8M Urea, 100mM Tris-HCl pH 8.0). Reduction was performed on top of the filters with 100 µL of 10mM TCEP in 8 M urea, 100mM Tris-HCl pH 8.0 at 37°C for 60 minutes with gentle shaking. Reduction solution was removed by extended centrifugation followed by two extra washing steps. Alkylation was then performed on top of the filters with 100 µL of 40mM chloroacetamide in 8 M urea, 100mM Tris-HCL pH 8.0 at 37°C for 45 minutes light protected with gentle shaking. After removal of alkylation solution and two additional washing steps with 200 µL of Urea solution, the filters were further washed twice with 200 µL of 5mM Tris-HCl pH 8.0. Samples were then digested overnight on top of filters at 37°C with 100 µL of proteolytic solution at 1/50 w/w enzyme-to-protein ratio using a combination of mass spectrometry grade trypsin and LysC supplemented with 10mM CaCl2. Elution from filters of resulting peptides was achieved by extended centrifugation and two extra elutions were performed with 50 µL of 4% TFA. Samples were then desalted on SDB-RPS Empore StageTips using the standard protocol^47^ and dried by vacuum centrifugation. Dried peptides were reconstituted in 10 µL 100mM HEPES pH 8 and then, 5 µL of TMT solution (25 µg/µL in pure acetonitrile) was directly added. Labeling was achieved with the TMTproTM 18plex isobaric Mass Tagging Kit for 90 minutes at room temperature and reactions were then quenched with hydroxylamine to a final concentration of 0.4% (v/v) for 15 min.. In order to avoid final extra normalization, a minor fraction of TMT labeled samples were pooled at 1:1 ratio across all samples and a single LC-MS2 run was performed to ensure correct peptide mixing. Corrected mixing quantities of each TMT-labeled sample were calculated based on the control run and final combined samples were desalted using a 100 mg Sep-Pak tC18 cartridge according to provider recommendations and dried by vacuum centrifugation. Pooled samples were fractionated into 12 fractions by Agilent OFF-Gel 3100 system following the manufacturer’s instructions. The resulting fractions were desalted again using SDB-RPS Empore StageTips and dried by vacuum centrifugation. Each fraction was resuspended in 2% acetonitrile, 0.1% Formic acid and nano-flow separations were performed on an Ultimate 3000 RSLC UHPLC system online connected with an Orbitrap Fusion Lumos Tribrid Mass Spectrometer (ThermoFisher Scientific). Samples were first trapped on a capillary pre-column (Acclaim Pepmap C18, 3 µm-100 Å, 2 cm × 75 µm ID), followed by an analytical separation at 250 nl/min over a 150 min. biphasic gradients through a 50 cm long in-house packed capillary column (75 µm ID, ReproSil-Pur C18-AQ 1.9 µm silica beads, Dr. Maisch). The data was acquired through a Top Speed Data-Dependent acquisition mode using three seconds cycle time. First MS scans were acquired at a resolution of 120,000 (at 200 m/z), with the most intense parent ions selected and fragmented by High Energy collision Dissociation (HCD) with a Normalized Collision Energy (NCE) of 35% using an isolation window of 0.7 m/z. Fragmented ions scans were acquired with a resolution of 50,000 (at 200 m/z) and selected ions were then excluded for the following 60 seconds. Resulting Raw files were searched using multiple Database search engines (Sequest HT, Mascot, MS Amanda and MSFragger) in Proteome Discoverer (v. 2.5) against the Uniprot_Mouse_55286Sequences_LR2022_05 supplemented with classical MaxQuant contaminants. Enzyme specificity was set to trypsin with a maximum of two missed cleavages allowed. A 1% FDR cut-off was applied both at peptide and protein identification levels. For the database search, carbamidomethylation (C), TMTpro tags (K and Peptide N termini) were set as fixed modifications while oxidation (M) was set as a variable one. Resulting text files were processed through in-house written R scripts (version 4.1.2). A first normalisation step was applied based to the Sample Loading normalisation ^48^. Assuming that the total protein abundances were equal across the TMT channels, the reporter ion intensities of all spectra were summed and each channel was scaled according to this sum, so that the sum of reporter ion signals per channel equals the average of the signals across samples. A second normalization was applied using TMM (trimmed mean of M-values), and dispersion estimates were obtained using a generalized linear model (GLM) framework. Log_2_ fold changes and p-values were calculated using quasi-likelihood F-tests, with multiple testing correction applied using the Benjamini-Hochberg (BH) method. The final dataset includes log_2_FC values and adjusted p-values for each condition compared to the control. All conditions were analyzed in triplicates, except for the Double condition, which was analyzed in duplicates.

### Protein degradation analysis and half-life correlation

To analyze protein degradation rates and their correlation with differential expression, protein degradation rate data were obtained from an external dataset ^49^, and compared to our proteomics dataset (see above). Differentially expressed proteins were identified for all starvation conditions based on significance thresholds (p < 0.01) and |log_2_FC| cutoffs (> 0.26). One-way ANOVA followed by Tukey′s post hoc test was conducted to determine statistical differences in half-life distributions between up and downregulated protein groups. Theoretical log_2_FC values were estimated based on protein half-life. A model was used to approximate the expected log_2_FC given a specific half-life, assuming a standard degradation kinetics equation (−6/half-life). Proteins with available degradation data were mapped to their theoretical log_2_FC, and significant downregulated proteins were highlighted. The correlation between theoretical and measured log_2_FC was computed to assess the agreement between expected and observed expression changes.

### tRNA charging assay

The protocol was adapted from ^50^. Cells were lysed in 500 µl TRIzol (Invitrogen) for 5 minutes at room temperature. After lysis, 100 µl chloroform was added, and samples were incubated on ice for 3 minutes. Following centrifugation (18,600 × g, 10 minutes, 4°C), the aqueous phase was transferred to a new tube containing 250 µl isopropanol, mixed, and incubated at −20°C for 10 minutes. RNA was pelleted (18,600 × g, 15 minutes, 4°C), washed with 80% cold ethanol, air-dried, and resuspended in 10 µL tRNA resuspension buffer (10 mM sodium acetate pH 4.5, 1 mM EDTA). For oxidation, 2 µg RNA was diluted in tRNA resuspension buffer (16.87 µL total volume) and incubated with 1 µL 0.2 M freshly prepared sodium periodate (NaIO₄) at room temperature for 20 minutes (control samples received 1 µL 0.2 M NaCl instead). The reaction was quenched with 2.75 µL 2.5 M glucose. After quenching, 1 µL E. coli spike RNA (4 ng/µL; Supplementary Material and Methods), 1 µL GlycoBlue (Invitrogen), and 1.3 µL 1 M NaCl were added. RNA was precipitated with 75 µL ice-cold 100% ethanol at −20°C for 30 minutes, pelleted (18,600 × g, 15 minutes, 4°C), washed with 70% ethanol, air-dried, and resuspended in 100 µL 50 mM Tris-HCl pH 9.0. Deacylation was performed at 37°C for 45 minutes, quenched with 100 µL tRNA quench buffer (50 mM sodium acetate pH 4.5, 100 mM NaCl), and precipitated overnight at −20°C with 540 µL ice-cold 100% ethanol. The RNA pellet was washed with 80% ethanol and dissolved in 5 µL water. For adaptor ligation, 800 ng RNA was mixed with 0.5 µL 10 µM tRNA adaptor (Supplementary Material and Methods), 1 µL 10x T4 RNA ligase buffer (NEB), 0.4 µL 0.1 M DTT, 2.5 µL 50% PEG, 0.5 µL murine RNase inhibitor (NEB), and 0.25 µL T4 RNA Ligase 2, truncated KQ (NEB) in a final volume of 10 µL. The reaction was incubated at room temperature for 3 hours, followed by overnight incubation at 4°C. Reverse transcription was initiated by annealing 0.5 µL 5 µM RT primer (Supplementary Material and Methods) to 2 µL ligated RNA (65°C for 5 minutes, then immediately chilled on ice). A master mix containing 0.8 µL 5x SuperScript IV buffer (Invitrogen), 0.2 µL 10 mM dNTPs, 0.2 µL 0.1 M DTT, 0.2 µL murine RNase inhibitor, and 0.2 µL SuperScript IV enzyme (Invitrogen) was prepared, and 1.6 µL was added to the annealed RNA (final reaction volume: 4 µL). Reverse transcription was performed at 55°C for 30 minutes, followed by inactivation at 80°C for 10 minutes. The cDNA was diluted to 15 µL with water. qPCR was performed using tRNA-specific primers (Supplementary Material and Methods), and Ct values were normalized to the E. coli spike-in control. Experimental samples under starvation conditions were normalized to control samples grown in full medium. For inhibition studies, cells were treated during starvation with either Bafilomycin A1 (160 nM, Sigma Aldrich) and MG132 (10 µM; Sigma Aldrich) to inhibit amino acid recycling, or cycloheximide (100 µg/mL; Sigma Aldrich) for the last 30 minutes to block translation.

### Analysis of transcripts involved in mTORC1 and GCN2 pathways

To analyze the activation of specific stress response pathways, we utilized the log_2_FC datasets of the Ribo-seq and RNA-seq. The ATF4 target genes were identified as the top 150 targets from ChIP-Atlas ^51^, while the mTORC1-sensitive genes were selected as known TOP transcripts from Thoreen et al ^22^. Statistical significance between conditions was assessed using unpaired t-tests.

### Preparation of genome-wide codon count table

The CDSs of the mouse genome (Mus musculus, GRCm39) were extracted and processed using R. Codon frequencies were extracted by splitting sequences into triplets. Codon counts were calculated for each gene, and isoform-specific counts were averaged to obtain gene-level codon frequencies. The codon count table was normalized by dividing each codon count by the total codon count per gene, resulting in the fraction of codons per gene. The genes included in this table were further filtered for transcripts detected to be expressed in mouse NIH3T3 cells according to our RNA-seq and Ribo-seq experiments. Thus, this table includes the relative usage of each codon across all expressed genes in the mouse NIH3T3 cell genome and was used for the downstream analyses.

### Processing & analysis of position-specific Ribo-seq data

Raw count data were filtered and normalized to the library size (to obtain positional RPM) and further normalized by dividing each RPM by the respective total RPM of the transcript (Norm_RPM). Genes with low counts (mean total counts across all samples below 50) were excluded. Coverage filtering was applied by calculating the fraction of positions with non-zero counts for each gene. Genes with mean positional coverage below 30% were excluded, resulting in a set of 4123 transcripts.

For codon-specific ribosome density enrichments, codons of interest were identified within the CDS, and positional Ribo-seq data were extracted in a window of ±50 nucleotides around each codon of interest. For each position, the relative Norm_RPM was computed as the ratio of experimental Norm_RPM to the control Norm_RPM baseline. Mean relative values across replicates were summarized for each codon and position.

For metagene profiles, positional RPMs were aligned relative to the start and end of the CDS. The normalized position was computed as a percentage of CDS length. Aggregated norm_RPM values were calculated across all transcripts for each condition and normalized position. To quantify ribosome accumulation at the 5′ and 3′ ends of transcripts, the summed norm_RPMs were calculated for three regions: the first 20% (5′ region), middle 20% (mid-region), and last 20% (3′ region) of the CDS. These scores were then normalized to the control condition by calculating the ratio of each condition score to the mean control score per gene.

### Polarity score analysis

The polarity score was used to quantify the bias of ribosome density between the 5′ and 3′ regions of CDSs and calculated as previously described ^52^. Conceptually, positions within the CDS that are closer to the 5′ end will result in a negative contribution to the polarity score, while positions closer to the 3′ end will contribute positively. This means that a higher polarity score indicates greater ribosome density near the 3′ end of the CDS, while a lower or negative polarity score suggests more ribosomes near the 5′ end. The Δpolarity score was calculated for each transcript by subtracting the Ctrl score and significance assessed by unpaired t-test. Transcripts with a significant increase in ribosome occupancy in the 5′ region in comparison to the Ctrl condition (Δpolarity < −0.15, p < 0.05) were extracted for further codon usage analysis. Codon frequencies were calculated for leucine, isoleucine, and valine codons in the first 20% of the CDS and normalized to the total codon count in that gene. Statistical comparisons were performed using unpaired BH-adjusted t-tests. The log_2_FC of codon usage was calculated relative to the control.

### Peak calling analysis

To identify positions and transcripts exhibiting ribosome stalling upon starvation, we computed for every transcript position the RPM and norm_RPM fold change relative to the control condition. Each position was also tested for having a higher RPM than the transcript mean. Statistical significance was assessed using unpaired t-tests. Peaks were defined based on the following criteria: log_2_FC(RPM) > 2, log_2_FC(Norm_RPM) > 2, and FC_relative_to_gene_mean > 1. Peaks that met all three thresholds and showed statistically significant differences (p < 0.05) between conditions were considered robust. To investigate the distribution of stalling sites along CDS, positions of upregulated stalling sites were mapped onto relative CDS coordinates. The spatial relationship between stalling events was examined by computing the distance between the first (most 5′) occurrence of Val and Ile stalling events within the same gene.

## Results

### Translation elongation rates are modulated in a codon-specific, non-additive manner upon limitation for BCAAs

To systematically assess how BCAA limitation affects translation elongation, we performed Ribo-seq on NIH3T3 cells starved for BCAAs — leucine, isoleucine, and valine. We examined the effects of single amino acid starvations (-Leu, -Ile and -Val), as well as combinations, including a double starvation of leucine and isoleucine (hereafter referred to as “double”) and a starvation of leucine, isoleucine, and valine (“triple”), allowing us to identify potential non-additive effects. Our experiments focused on a 6-hour time window to capture early cellular responses and avoid long-term adaptations or cell death. Across these Ribo-seq experiments, footprint lengths peaked at 31 nucleotides and reads predominantly mapped to coding sequences (Supplementary Fig. 1A-B). To examine how the limitation of BCAAs alters elongation, we inferred the genome-wide DT changes for all 61 sense codons across the different starvation conditions using our Ribo-DT pipeline ^44^. Notably, under control conditions, the DTs in our NIH3T3 cells were highly correlated to those previously reported in mouse liver, indicating that our system robustly captures mammalian translation elongation and is suitable for controlled nutrient-deprivation studies ^36^ (Supplementary Fig. 1C). Moreover, we noticed that DT changes extend beyond the ribosomal A-site, including the P-site, E-site, and even further positions (Supplementary Fig. 2A), consistent with other studies on single amino acid starvation ^39^ (Supplementary Fig. 2B-C). Consequently, we focused on a ±3 codon region centered on the P-site rather than a single codon position.

Intriguingly, only two of the three isoleucine codons (AUU and AUC) showed increased DTs upon Ile starvation (p < 0.01), while just one leucine codon (CUU) exhibited a modest but significant DT increase (p < 0.01) under Leu starvation (Figure 1A-B, Supplementary Figure 2A). In contrast, all four valine codons exhibited significantly increased DTs upon Val starvation (p < 0.01; Figure 1C). The DT changes were milder under double starvation, with only AUU (Ile) showing a significant DT increase among the BCAA codons (p < 0.01; Figure 1D). Surprisingly, under triple starvation, only valine codons exhibited significant DT increases (p < 0.01), with no significant changes observed for any of the leucine or isoleucine codons (Figure 1E). We complemented these observations by examining the averaged Ribo-seq read density around each codon corresponding to the starved amino acid. Under Val starvation, ribosomes strongly accumulated around all valine codons, whereas Leu starvation produced only minor peaks near CUU, CUC, and UUG, which diminished further during double or triple starvation (Figure 1F). Ile starvation caused a strong enrichment of ribosome density around AUC and AUU codons (Figure 1F). Notably, these enrichments persisted under double starvation but disappeared under triple starvation. These starvation-specific DT and ribosome density modulations were also evident at the individual transcript level, as exemplified by *Col1a1*, *Col1a2*, *Aars*, and *Mki67* which showed persistent Val-codon-specific ribosome density increases but lost Ile-codon-specific increases under triple starvation (Supplementary Figure 3A-D).

**Figure 1.**
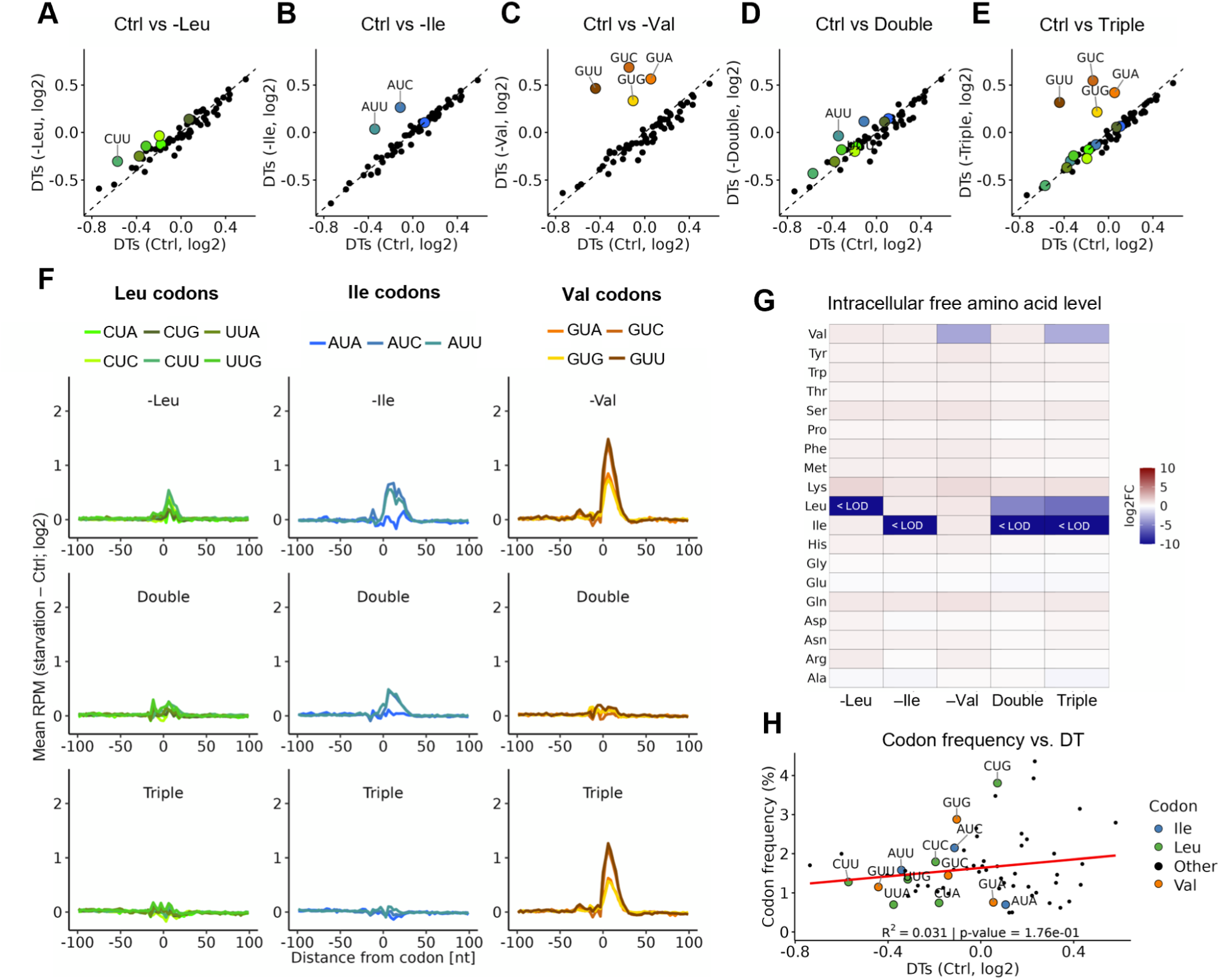
Translation elongation rates are modulated in a codon-specific, non-additive manner upon limitation for BCAAs. (A-E) Codon-specific ribosomal dwell times (DTs) in Ctrl vs. starvation (Double: -Leu & -Ile, Triple: -Leu & -Ile & -Val). Shown is the mean DT of triplicates over a ±3 codon region centered on the P-site. Significance was evaluated with ANOVA followed by Tukey’s HSD test. Annotated are codons with significantly upregulated DT relative to Ctrl (p-value < 0.01). (F) Changes in Ribo-seq read density around annotated codons relative to the Ctrl. Mean Reads per million (RPM) values were normalized to Ctrl condition, and the mean log_2_FC computed for a ±100 nt window around codons. (G) Heatmap of log_2_FC in intracellular amino acid levels under different starvation conditions relative to Ctrl. Values below detection limit (LOD) are indicated. (H) Scatter plot of DT under Ctrl condition (A-E, log₂) vs. codon frequency (%). Codons for Val (orange), Ile (blue), and Leu (green) are highlighted. A linear regression (red line) shows the relationship between DT and codon frequency (R² and p-value displayed).

Especially the observation that increased DTs were seen exclusively on valine codons under triple amino acid starvation was highly unexpected. To gain further insight into why valine codons uniquely displayed increased DTs under triple starvation and to exclude the possibility of compensatory mechanisms, we measured free intracellular amino acid levels. The deprived amino acids were drastically reduced or even undetected in their respective starvation conditions (Figure 1G), confirming effective cellular starvation. We next investigated whether differential codon usage frequency explained the observed DT changes, however, codons showing prolonged DTs were not particularly rare in transcripts expressed in mouse NIH3T3 cells (Supplementary Figure 3E). Moreover, there was neither a general correlation between DTs and average codon frequency (R^2^ = 0.031, p = 0.176) nor a bias in the effects of starvation on the DTs toward codons considered “slow” or “fast” (Figure 1H). Thus, the codon-specific stalling under BCAA starvation appears to be not driven by overall codon usage.

### Stress response pathways are differentially activated upon leucine, isoleucine and valine starvation

Prior work suggests that the activation of stress response pathways, particularly the GCN2 and mTORC1 pathways, influence ribosome stalling during amino acid starvation ^39^. In particular, a strong activation of these stress pathways (e.g., under leucine deprivation) prevented ribosome stalling by efficiently reducing global translation load, whereas weaker modulations (e.g., arginine deprivation) led to more frequent stalling. Therefore, we evaluated the activation of stress pathways across our BCAA starvations.

Our RNA-seq and Ribo-seq revealed a general activation of stress response pathways across all starvations. Principal Component Analysis (PCA) of mRNA levels, ribosome footprints (RPFs), and ribosome densities (RD = Ribo-seq/RNA-seq) showed a certain level of shared transcriptional and translational reprogramming across starvations (Supplementary Figure 4A-B; PC1), involving pathways related to the unfolded protein response (UPR), ubiquitin-like protein binding, and cell cycle regulation (sister chromatid segregation) (Supplementary Figure 4D-E, Supplementary Data 3), fundamentally interconnected processes in the cellular stress response. Notably, Val and triple starvation clustered close together - further from the control than the other conditions - especially at the ribosome footprint level, suggesting a stronger translational impact (Supplementary Figure 4B). Interestingly, Ile starvation exhibited a distinct pattern, especially evident in the Ribo-seq (Supplementary Figure 4B), and this distinction became even clearer at the RD level (Supplementary Figure 4C), where Ile starvation separated from the Ctrl only along PC2. Transcripts driving this separation were significantly enriched for genes involved in the regulation of transcription in response to stress and cytoplasmic translation (Supplementary Figure 4F, Supplementary Data 3). Consistent with that, pairwise comparisons of RNA-seq, Ribo-seq and RD changes (often used as a proxy for changes in translation efficiency) showed that Val starvation caused the strongest changes, with 423 mRNAs showing increased RD and 403 mRNAs showing decreased RD (Supplementary Figure 5A, Supplementary Data 3). These changes largely overlapped with those observed under triple starvation, whereas Ile starvation selectively impacted a unique subset of transcripts, many of which linked to cytoplasmic ribosomal proteins (Supplementary Figure 5B, D). Transcripts showing an altered RD under all BCAA starvations included key translation initiation factors, such as *Eif4g3* and *Eif1*, several transcription factors, as well as UPR-associated genes, such as *Chop*, *Atf4*, *Ppp1r15a* and *Yod1* (Supplementary Figure 5B-C).

CHOP and ATF4 are stress-induced transcription factors that mediate the GCN2 pathway. We observed a strong upregulation of *Atf4* and *Chop* at both the mRNA and ribosome footprint levels (Figure 2A) across all starvations, indicating a robust activation of the ISR. This activation was further validated by qPCR profiling of the transcriptional activation of ISR genes (*Asns*, *Atg3*, and *p62*) over a 3- to 6-hour time course, with most genes showing a robust induction after 6h of all starvations (Figure 2B). Of note, double starvation resulted in the weakest transcriptional activation of the tested ISR genes and a lower induction of *Chop* expression (Figure 2A-B). Nevertheless, RNA-seq analysis of direct ATF4 target transcripts confirmed their significant upregulation, without significant differences between the starvation conditions (Figure 2C). To assess the mTORC1 branch, we measured RPS6 and 4E-BP1 phosphorylation by Western blot. Notably, double starvation, and to a certain extent individual Leu and Ile starvation, decreased RPS6 phosphorylation, indicating robust mTORC1 inhibition, while Val and triple starvation resulted in milder changes (Figure 2D-E). A similar trend was found for 4E-BP1, albeit less pronounced (Figure 2D-E). In line with this, ribosome coverage of mTORC1-sensitive TOP transcripts was typically lowered upon Leu and double starvation but not as consistently altered upon Val or triple starvation (Figure 2F). Because TOP transcripts predominantly encode ribosomal proteins (RPs), which are prone to multimapped reads, it can be challenging to accurately quantify them in Ribo-seq data. Therefore, we repeated the analysis and included multimapped reads to more reliably assess ribosome coverage (Supplementary Figure 5E). This analysis confirmed the original trends and revealed even more pronounced downregulation, especially under double starvation, and slightly less pronounced but still evident reductions upon Leu and Ile starvation.

**Figure 2.**
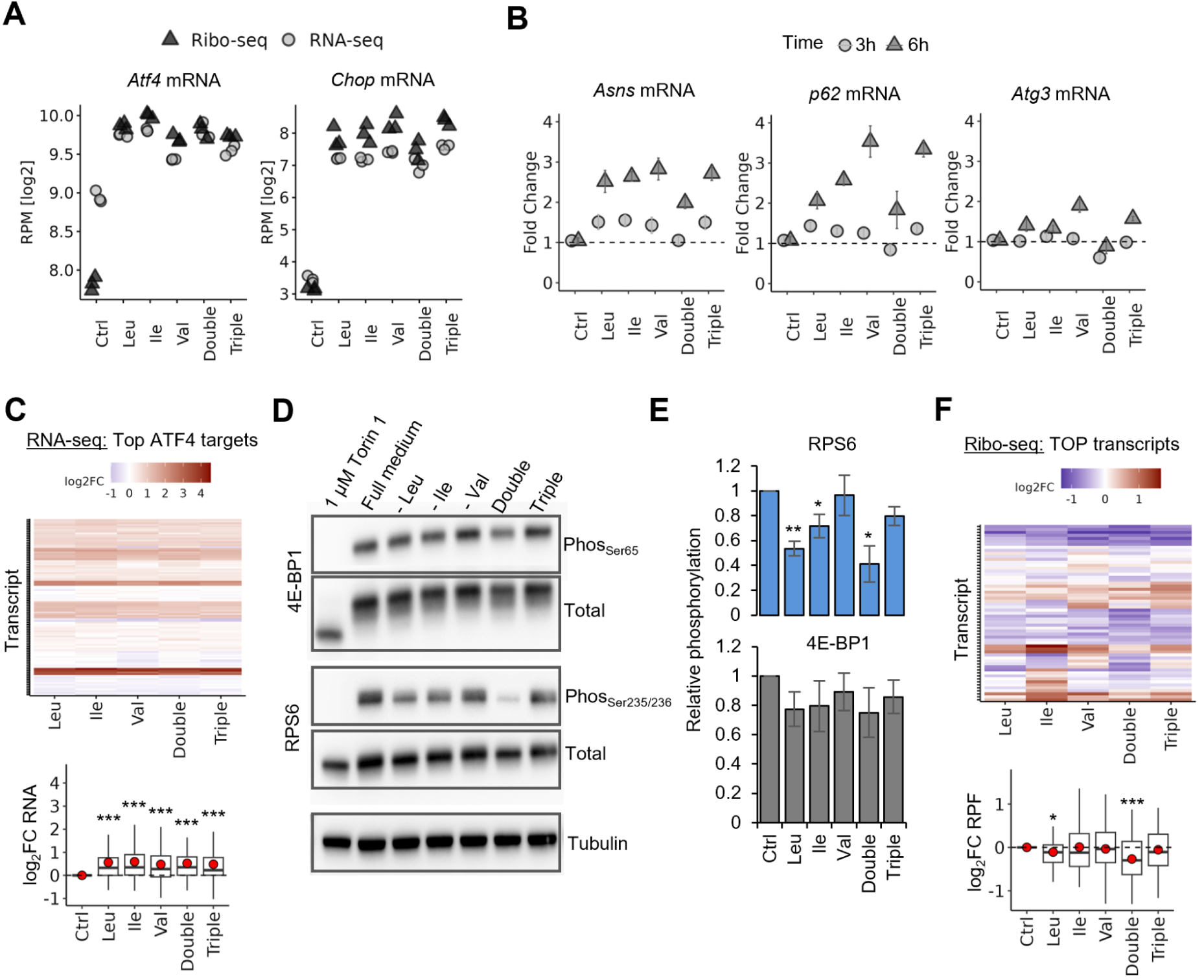
Stress response pathways are differentially activated upon Leu, Ile and Val starvation. (A) Expression levels of *Atf4* and *Chop* in RNA-seq and Ribo-seq under different deprivation conditions. Values represent log₂-transformed reads per million (RPM). Each point represents an individual replicate. Statistical significance was determined by unpaired t-tests comparing each starvation to Ctrl, with all comparisons resulting in p < 0.01. (B) qPCR analysis of mRNA levels for *Asns*, *p62*, and *Atg3* under different starvations at 3h and 6h. Expression levels were normalized using the ΔΔCt method (normalized to *Gapdh* expression) and are shown as fold change relative to Ctrl. Bars represent mean fold change, with error bars indicating standard error of mean (SEM). (C) Heatmap and quantification of log₂FC for the top 150 ATF4 target genes from ChIP-Atlas^51^ at mRNA level. Genes are clustered based on their expression profiles using k-means clustering (k = 12). In the boxplot, red points indicating the mean expression change. Significance was assessed using unpaired t-tests. (D) Representative Western blot showing phosphorylation levels of 4E-BP1 (Phos_Ser65_) and RPS6 (Phos_Ser235/236_) under different amino acid starvations. Total protein levels and Tubulin level serve as loading controls. 1 µM Torin1 treatment and full medium conditions are included as controls. (E) Quantification of phosphorylation levels of 4E-BP1 and RPS6, normalized to total protein levels and relative to the control (Ctrl). Quantification was performed using ImageJ. Data represent mean ± SEM from three independent experiments. Statistical significance was determined using unpaired t-tests. (F) Heatmap and quantification of log₂FC for the known TOP motif containing transcripts^22^ in our Ribo-seq. Genes are clustered based on their expression profiles using k-means clustering (k = 12). In the boxplot, red points indicating the mean expression change. Significance was assessed using unpaired t-tests.

Thus, while all starvations triggered stress responses, the exact balance of GCN2 and mTORC1 pathway engagement differed depending on which BCAAs were depleted. This differential pathway activation may play a role in why the extent of effects observed on the ribosome DTs is milder during double and Leu starvation.

### BCAA starvations differentially modulate translation initiation and elongation

Because both the modulation of the GCN2 and mTORC1 pathway aim to decrease translation initiation, we examined to what extent overall protein synthesis rates were reduced under each starvation condition. We treated cells with a short pulse of OPP (a puromycin alkyne analog) after the starvation periods, then used click-chemistry to fluorescently tag newly synthesized proteins with TAMRA-azide for quantification ^53^ (Figure 3A). Cells treated with cycloheximide (CHX), a translation inhibitor, and OPP-untreated cells confirmed that the observed fluorescence signal was due to active translation (Figure 3B-C). In all starvation conditions, TAMRA signal intensity was significantly reduced, confirming a decrease in global protein synthesis rates (Leu: 50%, Ile: 46%, Val: 48%, Double: 54%, Triple: 65%; Figure 3D). As a parallel approach to analyze global translation, we also performed polysome profiling (Figure 3E). We observed that starving cells for Leu, Val, and their combinations lead to significant increases in monosome levels and a decrease in polysomes, as evidenced by a reduced polysome-to-monosome (P/M) ratio of about 50% (Figure 3F-H), indicating globally fewer ribosomes per transcript likely as a consequence of reduced initiation. By contrast, the starvation of Ile alone did not alter the P/M ratio or the levels of polysomes (Figure 3F,H). Importantly, reduced translation elongation can have an opposite effect than reduced initiation on the amount of measured polysome. Elongation slowdown can result in accumulation of ribosomes on transcripts, thereby increasing the amount of polysomes. Under starvation, both initiation and elongation are affected and the observed polysome profiles reflect a blend of these opposing effects. Therefore, the unaltered P/M ratio upon Ile starvation, in combination with an reduced OPP incorporation points towards both reduced translation initiation and globally slower elongation, potentially corroborating the distinct translational signature of Ile starvation in the Ribo-seq data (Supplementary Figure 4A-C). Together, these findings highlight that BCAA starvation triggers a combination of effects on initiation and elongation, with varying dynamics by amino acid starvation.

**Figure 3.**
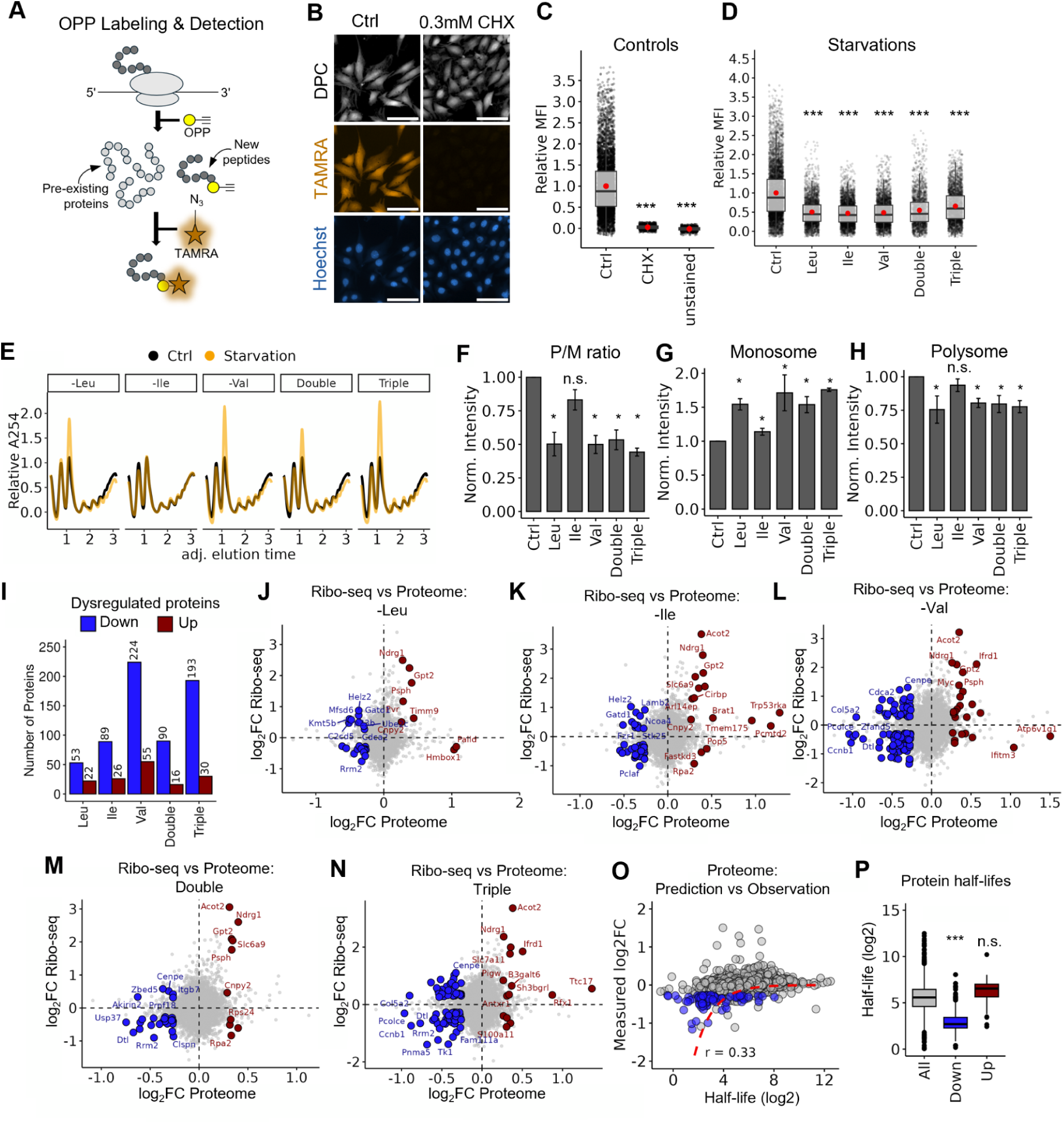
BCAA starvations differentially modulate translation initiation and elongation. (A) Schematic representation of detection of newly synthesised proteins by OPP incorporation assay. Newly synthesized proteins incorporate OPP, which causes translation termination and allows subsequent labeling with TAMRA-azide. This enables their detection and quantification via fluorescence microscopy, distinguishing them from pre-existing proteins that lack OPP. (B) Representative fluorescence microscopy images of TAMRA and Hoechst staining. Wild-type cells and cells treated with 0.3 mM CHX to inhibit translation are shown. TAMRA (orange) represents OPP staining, marking newly synthesized proteins. Hoechst stains nuclei (blue), and DPC (digital phase contrast) provides cellular morphology. Scale bar: 100 µm. (C-D) Quantification of OPP incorporation under (C) Ctrl and (D) starvation conditions. Boxplot representations display the relative mean fluorescence intensity (MFI) of the TAMRA signal; each dot represents an individual cell, and red points indicate the mean MFI. Data represent three biological replicates. Statistical significance between conditions was determined using unpaired t-tests with Bonferroni correction. (E) Representative polysome profiles of Ctrl (black line) and starvations (orange line). (F–H) Quantification of polysome-to-monosome (P/M) ratio (F), monosome (G), and polysome (H) fractions. (I) Number of dysregulated proteins (p < 0.01 and |log₂FC| > 0.26) across starvations. (J–N) Scatter plots showing the relationship between log_2_FC measured in proteome and Ribo-seq under the given conditions. Red dots represent transcripts corresponding to upregulated proteins (p < 0.01 and log₂FC > 0.26) with significant changes in Ribo-seq (p < 0.01 and |log₂FC| > 0.26), while blue dots highlight transcripts corresponding to downregulated proteins (p < 0.01 and log₂FC < −0.26) with significant changes in Ribo-seq (p < 0.01 and |log₂FC| > 0.26), indicating translational repression. (O) Correlation between measured log₂FC proteomic changes upon Val starvation and published protein half-lifes in wiltype NIH3T3 cells ^49^. Red line indicates theoretical log_2_FC assuming standard degradation kinetics under total synthesis block. (P) Distribution of protein half-lives (see O) for down- and upregulated proteins under any BCAA starvation.

To determine whether these global changes in protein synthesis after 6 h of starvation are already reflected at the proteome level, we performed TMT-based quantitative proteomics. Of note, given our relatively short 6 h period of amino acid starvation, the changes in protein abundance are expected to be of limited magnitude. Among 7,004 proteins identified (6,177 also covered in RNA-seq and Ribo-seq; Supplementary data 3), Val and triple starvation caused the largest proteome changes (224/193 downregulated proteins, respectively; Figure 3I), whereas Leu starvation had the mildest effect on the proteome (53 downregulated, 22 upregulated; Figure 3I). Interestingly, while many of the downregulated proteins showed a decreased level of ribosome footprints (Figure 3J-N; blue dots), and upregulated proteins showed an increase in the footprint level (Figure 3J-N; red dots), aligning with the hypothesis of global translation initiation modulation, we noticed that a subset of downregulated proteins maintained or even increased the ribosome occupancy of the corresponding transcript (Figure 3J-N; blue dots). Thus, for these proteins, the changes in protein abundance did not correlate with the ribosome footprint data, suggesting potential ribosome stalling (in line with our observations of DT changes in Figure 1). Such stalling could impair effective protein synthesis despite sustained or even increased ribosome occupancy.

Because protein stability modulated by protein degradation pathways can also alter protein levels, we compared the measured proteomic fold changes with published half-lives of these proteins ^49^. Downregulated proteins generally had shorter half-lives (Figure 3O-P), which likely made them more prone to rapid depletion once their synthesis was reduced. In addition, we modelled the theoretical log_2_FC values based on protein half-lifes, assuming standard first order degradation kinetics. We observed that measured protein fold changes were often less pronounced than predicted if translation had halted completely (Figure 3O; prediction indicated by red line), suggesting that increased proteolysis is unlikely to be the sole driver and the reduced protein level may also result from decreased or halted translation. While these comparisons cannot rule out the influence from protein stability changes, these results imply that BCAA deprivation lowers protein output through multiple pathways: a combination of reduced initiation, direct elongation blocks (stalling), and possibly an increased proteolysis.

### BCAA starvation leads to ribosome density shifts towards the 5′ of the CDS

While our global measurements established that BCAA starvation reduces protein synthesis and our Ribo-seq data suggests the involvement of ribosome stalling, the analysis did not indicate the regions along the mRNAs where the stalling happens. Therefore, we next performed metagene analysis of ribosome footprints, focusing on whether BCAA deprivation leads to shifts in ribosome density along the coding sequence. Of note, the first/last 15 nt of each transcript were excluded from the analysis to avoid distinct effects associated with the start/stop codon peaks ^54^. Strikingly, under all BCAA starvations, we detected ribosome accumulations at the 5′ ends of transcripts and depletions in their 3′ regions (Figure 4A). To quantify this effect, we calculated the transcript-specific differential ribosome occupancy ratios in the first 20% (5′ index) vs. the last 20% (3′ index) of CDSs, confirming a significant shift toward 5′-end accumulation under all starvations (Figure 4B).

**Figure 4.**
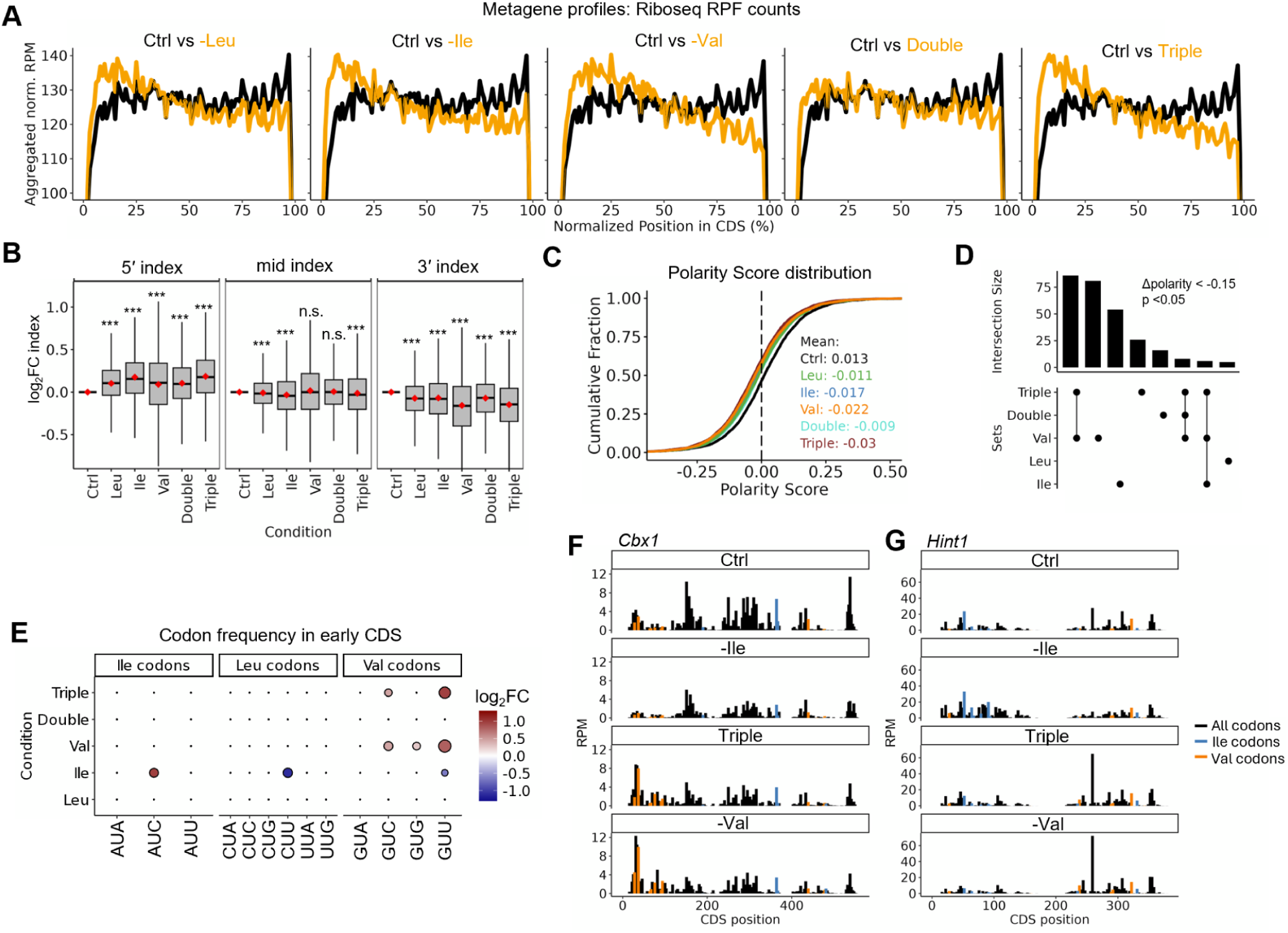
BCAA starvation leads to ribosome density shifts towards the 5′ of the CDS. (A) Metagene profiles showing the aggregated normalized RPF count of the Ribo-seq along the CDS for each starvation (orange) compared to Ctrl (black). Solid lines represent the mean aggregated signal at each normalized CDS position, and shaded ribbons indicate the standard error of the mean (SEM). (B) Boxplots quantifying the regional “ramp” in ribosome densities. Log₂-transformed ratios of summed RPF counts in the 5′ region (1% - 20% of CDS positions), mid-CDS (40% - 60% of CDS positions), and 3′ regions (80% - 100% of CDS positions) are shown for starvation relative to Ctrl. Red dot indicates the mean value. Statistical significance between Ctrl and starvation was determined using unpaired t-tests with Bonferroni correction. (C) Cumulative fraction plots of transcript-specific polarity scores. Mean polarity scores per condition are shown for each condition. (D) Overlap plot depicting the intersections of transcripts with increased 5′ coverage (Δpi < - 0.15, p < 0.05) across conditions. The bar plot represents the size of each intersection. (E) Dot plot summarizing changes in mean codon frequencies of transcripts with polarity score shift upon annotated starvations in the first 20% of their CDS. For each codon group (Val, Leu, and Ile codons), the log₂FC relative to remaining transcripts is plotted. Dot size reflects the –log₁₀(p-value) from unpaired t-tests (with BH adjustment), and the fill color (ranging from dark blue to dark red) indicates the magnitude and direction of change. (F–G) Representative ribosome profiling tracks for the genes *Cbx1* (F) and *Hint1* (G), with Val and Ile codons annotated.

To identify the most affected transcripts, we calculated polarity scores that reflect the distribution of ribosome density from 5′ to 3′. Although all BCAA limitations slightly shifted these polarity scores, Val and triple starvation produced the largest changes (Figure 4C; Supplementary Figure 6A). Notably, transcripts with significantly reduced differential polarity scores (Δpolarity < −0.15; p < 0.05) compared to Ctrl largely overlap between Val and triple conditions, while Ile starvation affected a mostly distinct set of transcripts (Figure 4D). Leu and double starvation resulted only in a small set of transcripts with a significant shift in polarity score (Figure 4D). Strikingly, for transcripts with reduced polarity scores under Ile or Val starvation, the 5′ CDS region contained a significantly higher frequency of codons cognate to the limiting amino acid compared with the 5′ CDS region of transcripts that did not exhibit a polarity shift (Figure 4E). Under triple starvation, however, again only valine codons were enriched in the early CDS of transcripts with reduced polarity scores. For example, *Cbx1*, a transcript with a high polarity shift under Val and triple starvation, contains multiple valine codons in its 5′ region, precisely where the ribosomes piled up (Figure 4F). Similarly, *Hint1*, affected by Ile starvation, has isoleucine codons near its start codon in regions with increased ribosome accumulations (Figure 4G). Notably, this codon enrichment was specific to the 5′ regions of stalled transcripts; the overall CDS of these transcripts did not show elevated frequencies of the limiting amino acid′s codons (Supplementary Figure 6B). Overall, our results suggest that BCAA deprivation alters ribosome distribution along transcripts, with distinct codon- and condition-specific effects on elongation. The codon-specific effects indicate that elongation bottlenecks may arise when ribosomes encounter codons for limiting amino acids, leading to stalled progression and the accumulation of ribosomes in upstream regions.

### Non-uniformly distributed codons create elongation bottlenecks

Despite the 5′-enriched ribosome accumulation under Val and triple starvation, it remains puzzling why cells upon triple starvation fail to show Ile-starvation-specific stalling, which were observed under single Ile starvation. Moreover, the metagene profiles did not pinpoint the exact codons along individual transcripts at which ribosomes paused. To identify these sites more precisely, we next aimed to systematically map ribosome stalling events by peak-calling (see Material and Methods). Strikingly, this approach revealed strong differences in the frequency and distribution of stalling events across the different starvations: Val starvation caused the highest number of stalling events (11953 sites across 3197 genes), which exceeded those observed under Leu, Ile, or double starvation (Figure 5A; Supplementary Figure 7A).

**Figure 5.**
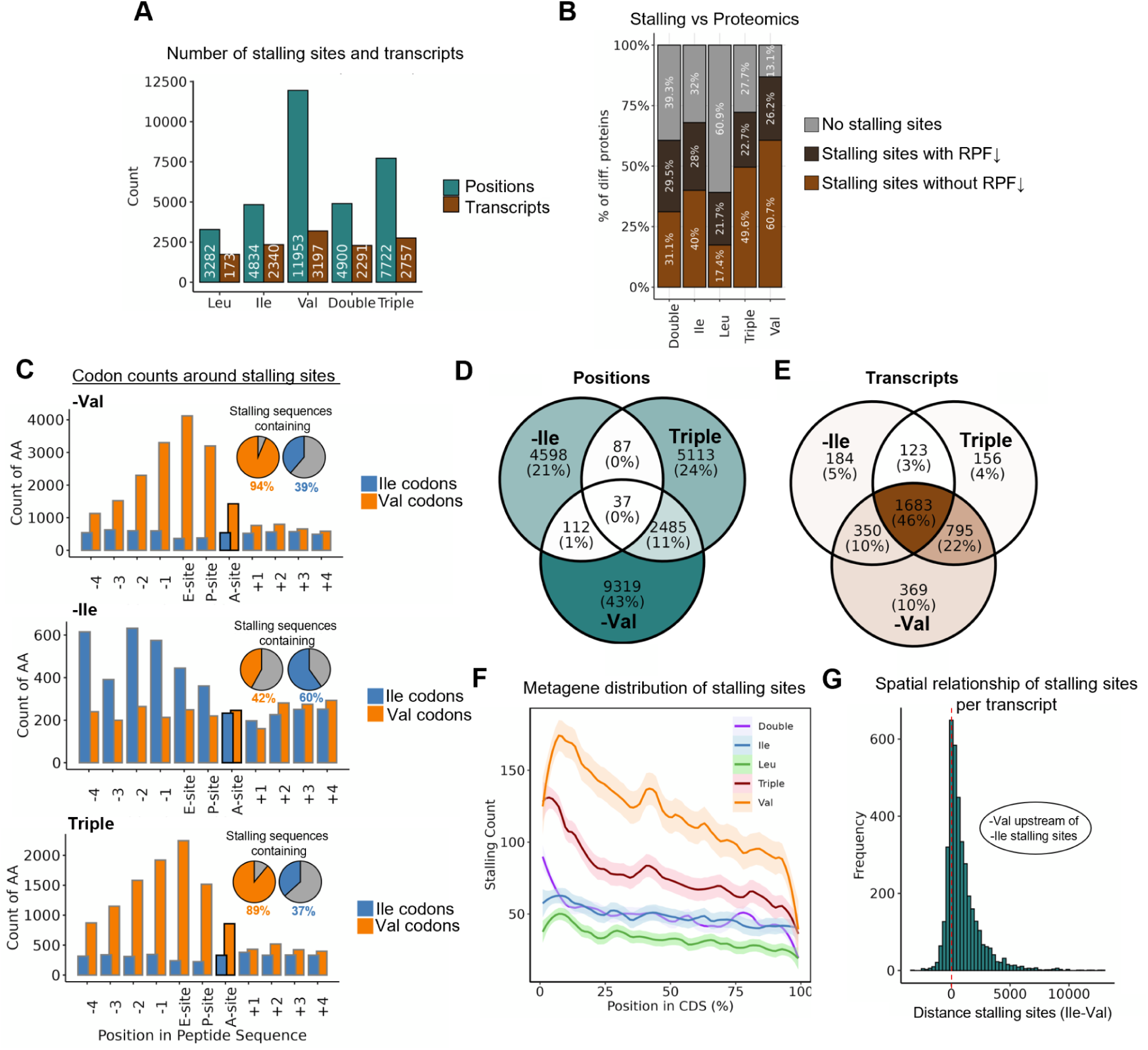
Non-uniformly distributed codons create elongation bottlenecks. (A) Bar plots showing the number of extracted stalling sites (positions) and the corresponding transcripts harboring stalling sites across different starvations. Peaks were defined based on the following criteria: log_2_FC(RPM) > 2 & log_2_FC(Norm_RPM) > 2 & FC_relative_to_gene_mean > 1. Peaks that met these thresholds and showed statistically significant differences (p < 0.05) between conditions were considered robust. (B) Comparison of downregulated proteins in each starvation to transcripts harbouring stalling sites (as identified in A). Gray bar indicates the percentage of downregulated proteins without stalling site and brown indicates proteins with an identified stalling site, with dark brown indicating an additionally downregulated RPF level (measured by Ribo-seq) and bright brown indicating no corresponding downregulated RPF level. (C) Count graphs depicting the valine and isoleucine codon counts at indicated positions around the extracted stalling sites in the annotated starvation condition. The region surrounding each stalling site was extracted and the amount of Val or Ile codons was summed at each position; accompanying pie charts indicate, in percentage, the proportion of extracted stalling sequences that contain an Ile or Val codon versus those that do not. (D-E) Venn diagrams displaying the overlap of identified (D) stalling positions and (E) transcripts that harbor stalling sites in Ile, Val, and Triple starvation. (F) Metagene distribution of the extracted stalling sites across the CDS in all starvation conditions. (G) Spatial analysis of stalling sites on the 1,683 transcripts harboring stalling sites in Ile, Triple, and Val starvation conditions. Per transcript, the distance between the first stalling site caused by -Ile starvation and the first stalling site caused by -Val starvation was calculated, revealing that under Val starvation the stalling occurs further upstream on the transcript.

Intriguingly, cross-referencing the transcripts with identified stalling sites with the proteomic data revealed that the majority of proteins downregulated under Val starvation (see Figure 3) harbored stalling sites (86.9%; hypergeometric p = 0.0233), while proteins downregulated under Leu starvation more rarely overlapped with them (39.1%; hypergeometric p = 0.750; Figure 5B). In addition, a high number of downregulated proteins with associated ribosome stalling sites did not show an overall decreased mean RPF count in ribosome profiling data (Figure 5B), as it would be expected from translation initiation defects, linking these stalling sites directly to proteomic changes.

To further address why the identified Ile starvation-specific stalling sites are not found upon triple starvation, we examined their local sequence context. Under Val and triple starvation, we found that valine codons were strongly enriched directly at or just upstream of the identified stalling positions (A-site), whereas isoleucine codons were scarce and uniformly distributed (Figure 5C). The pattern was reversed in cells upon Ile starvation, where isoleucine codons were abundant at or in close proximity to the stalling sites and valine codons were rarely seen (Figure 5C). Of note, in cases where valine or isoleucine codons were present just upstream (rather than at) the stalling position, we noted a strong bias for GAG (E), GAA (E), GAU (D), GAC (D), AAG (K), CAG (Q), GUG (V) and GGA (G) (Val starvation) and AAC (N), GAC (D), CUG (L), GAG (E), GCC (A), CAG (Q), GAA (E) and AAG (K) (Ile starvation) at the stalling site (Supplementary Figure 7B). Consistent with the previous analysis, Val and triple starvation showed a substantial overlap in their stalling sites, whereas few Ile-starvation-specific stalling positions persisted in triple starvation (Figure 5D). Nonetheless, when we examined entire transcripts rather than single positions, many transcripts that exhibited isoleucine-related stalling under Ile starvation also stalled under triple starvation, but at different sites along the CDS (Figure 5E). This finding is particularly intriguing, as it suggests that while Ile-starvation-specific stalling sites may shift under triple starvation, the overall tendency of these transcripts to stall remains.

Given this dynamic redistribution, we focused on the positional patterns of stalling sites along the CDS and observed that under Val and triple starvation, stalling events tended to cluster toward the 5′ end of the CDS, whereas in the other starvations, stalling sites appeared more evenly distributed (Figure 5F). Interestingly, these positional preferences mirrored the natural codon usage biases: We noticed a transcriptome-wide overrepresentation of valine codons in the early CDS, whereas isoleucine codons were comparatively more abundant toward the 3′ end (Supplementary Figure 7C-D). This observation gives rise to a potential explanation as to why triple starvation stalling rarely targeted isoleucine sites: Under triple starvation, an early stall of ribosomes on valine codons may create a bottleneck that may prevent or delays ribosomes from reaching downstream isoleucine codons (potentially owing to terminal stalling), thereby reducing the frequency of Ile-starvation-specific stalling. This concept was supported by detailed transcript-level analyses revealing that, in transcripts containing both Val- and Ile-starvation-specific stalling sites in their respective single starvation, the Val-starvation-specific stalling sites tend to lay upstream of the Ile-starvation-specific site (Figure 5G). Taken together, these observations demonstrate that BCAA limitation — in particular, Val starvation — establishes a prominent translation bottleneck early in elongation, ultimately impacting downstream ribosome occupancy and overall protein output.

### Differential tRNA charging patterns reveal a mechanistic basis for codon-specific stalling under BCAA starvation

Given our observations that early ribosome stalling under Val and triple starvation correlates with an overrepresentation of valine codons, we next investigated if the charging levels of tRNA isoacceptors under these conditions fully reflect the apparent elongation bottleneck. If valine tRNAs become depleted more quickly than the other tRNAs under the triple starvation, this could explain why ribosomes stall on valine-rich regions near the 5′ end of transcripts and fail to progress to downstream codons. Moreover, differences in the charging status of tRNA isoacceptors could underlie the codon-dependent stalling patterns observed under isoleucine starvation (Figure 1). To test this, we systematically measured the charging levels of tRNA isoacceptors for leucine, isoleucine, and valine (Figure 6A) in each starvation condition using periodate oxidation and tRNA isoacceptor-specific RT-qPCR (adapted from ^50^; see Material and Methods).

**Figure 6.**
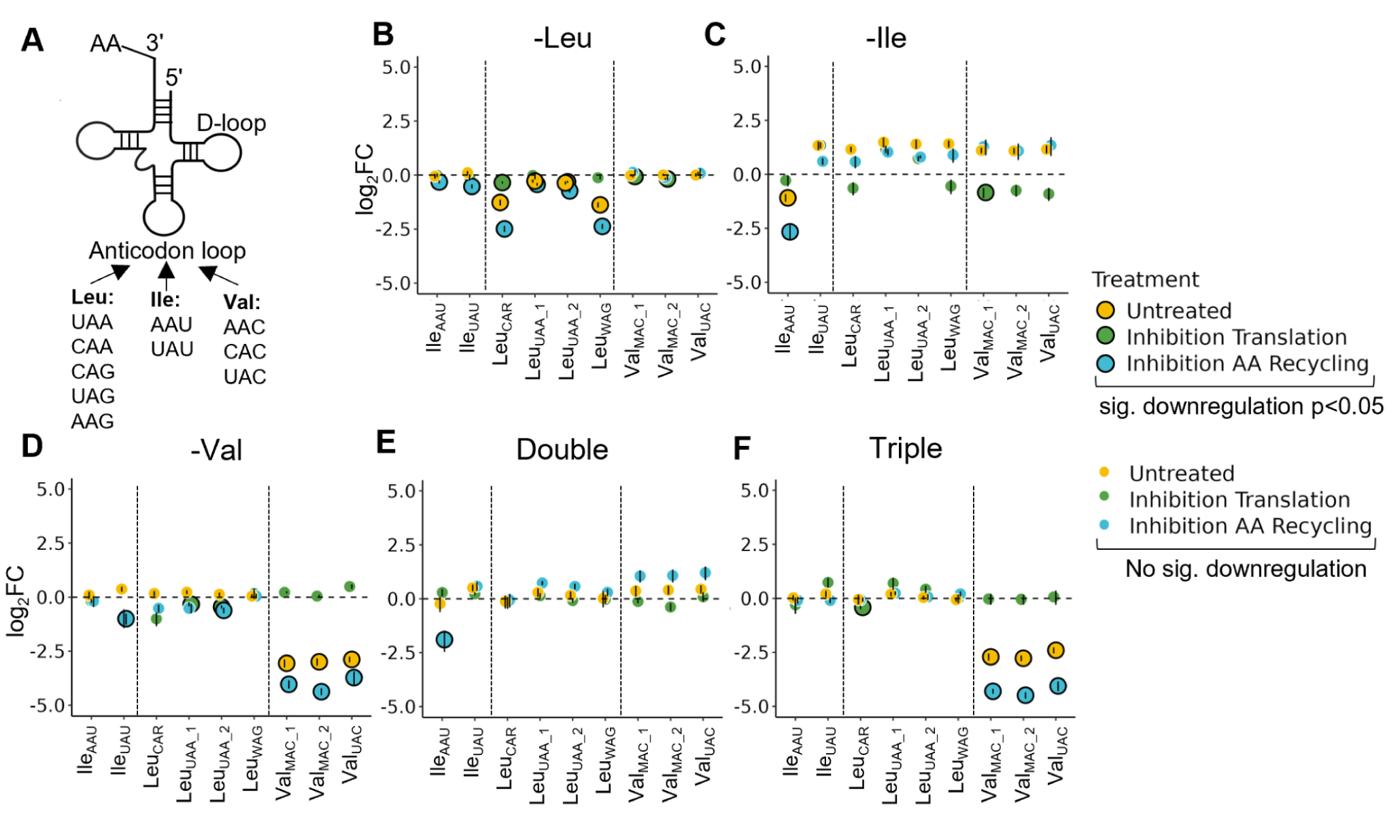
Differential tRNA charging patterns reveal a mechanistic basis for codon-specific stalling under BCAA starvation. (A) Schematic representation of the different tRNA isoacceptors for Leu, Ile, and Val. (B–F) Differential charging of tRNA isoacceptors under the indicated starvation conditions. Yellow indicates starved cells without treatment. Green represents cells treated with 100 µg/mL cycloheximide (CHX) in the last 30 min of starvation to inhibit translation. Blue represents cells treated with 10 µM MG132 and 160 nM Bafilomycin A1 to inhibit proteasomal and autophagic degradation (amino acid recycling inhibition). Certain tRNA isodecoders could not be measured separately and are therefore represented with IUPAC ambiguity codes: W (A or U), R (A or G), and M (A or C). Large outlined dots indicate significant downregulation (p < 0.05). Sample sizes: untreated (n = 5), translation and AA recycling inhibition (n = 2). Displayed is the mean of the replicates with the SEM indicated as error bars.

Aligning with our stalling data, Val and triple starvation both resulted in a significant decrease of all charged tRNA^Val^ isoacceptors (Figure 6D, F; yellow dots), while the triple starvation did not reduce charging for tRNA^Ile^ or tRNA^Leu^ (Figure 6F; yellow dots). By contrast, individual Leu starvation selectively reduced the charging of tRNA^Leu^ and tRNA^Leu^ (while sparing tRNA^Leu^) (Figure 6B; yellow dots). Similarly, upon Ile starvation, only tRNA^Ile^ lost charging, whereas tRNA^Ile^ did not (Figure 6C; yellow dots). Upon double starvation we could not observe a charging loss of any of the BCAA tRNAs (Figure 6E; yellow dots). These isoacceptor-specific patterns correlate largely with the particular subsets of leucine and isoleucine codons that stalled (Figure 1A). Of note, total tRNA levels remained largely unchanged across all starvation conditions (Supplementary Figure 8A-E), suggesting that observed differences in charged tRNAs reflect genuine shifts in aminoacylation rather than alterations in overall tRNA abundance.

To clarify whether active translation directly consumes these aminoacyl-tRNA pools leading to accumulation of uncharged tRNAs, we treated cells with CHX for the final 30 minutes of deprivation. CHX blocks ribosome translocation and thus ongoing elongation. Notably, CHX rescued the depletion of tRNA charging in each condition (Figure 6B-F; green dots), suggesting that, when the ribosomes are not actively translating, aminoacyl-tRNA synthetases can adequately charge tRNAs even in face of limited amino acids. Lastly, we probed whether amino acid recycling via autophagy and proteasomal degradation affected tRNA charging and would lead to the accumulation of the uncharged version of additional tRNA isoacceptors. Inhibiting these pathways upon triple starvation did not cause a loss of any tRNA^Ile^ isoacceptor charging, but it did intensify the charging loss of tRNA^Val^ isoacceptors (Figure 6F; blue dots). This aligns with our model that, upon triple starvation, ribosomes stall at valine codons near the transcript 5′ end, which can compromise their progression to isoleucine codons (e.g. in the event of terminal stalling at valine codons). As a consequence the demand for charged tRNA^Ile^ would be reduced and blocking autophagy and proteasomes would not exacerbate charged tRNA^Ile^ depletion. By contrast, charged tRNA^Val^ are used immediately and become more depleted when proteolytic recycling is cut off.

Thus, tRNA isoacceptor charging dynamics closely reflect the codon usage and ribosome stalling patterns observed under BCAA starvation. Our data strengthen the link between codon bias, localized tRNA shortages, and the translational blocks that arise under nutrient-limited conditions.

## Discussion

Cells dynamically regulate translation in response to nutrient availability, ensuring adaptive protein synthesis under stress conditions. In this study, we present a comprehensive analysis of how cells modulate translation in response to BCAA starvation. By combining data from RNA-seq, Ribo-seq, quantitative proteomics, and tRNA charging assays, we revealed codon-specific ribosome stalling events triggered by the starvation of leucine, isoleucine, and valine. These events are associated with altered tRNA charging levels, activation of stress response pathways, and show codon positional effects that influence ribosome stalling.

Our findings reveal that tRNA charging tightly correlates with the observed stalling patterns, providing a mechanistic basis for the codon-specific translational effects. Under Val starvation, all valine tRNA isoacceptors exhibit significantly reduced charging levels, whereas Ile and Leu starvation affect only specific tRNA isoacceptors. This observation aligns with early studies in *E. coli*, which predicted and later confirmed that when a single amino acid becomes limiting, certain tRNA isoacceptors lose their charging much faster than others, depending on their abundance and the frequency of the codons they read ^31,32^. Moreover, the selective tRNA charging under stress conditions has been shown in mammalian cells to modulate translation rates between stress-responsive and growth-associated transcripts ^55^. Importantly, our data indicate that a specific tRNA isoacceptor does not always become uncharged when its cognate amino acid is starved (i.e. seen in the triple starvation). In fact, even under complete amino acid starvation, only a subset of tRNAs lose charging ^50^, suggesting that tRNA charging dynamics are governed by a balance between supply and demand. This redistribution of available translational resources allows ribosomes and tRNAs to be repurposed for the translation of stress-response transcripts ^31,32^. For instance, in our double starvation condition, unchanged tRNA charging levels (Figure 6E) may result from a pronounced downregulation of global translation initiation, likely driven by the activation of stress responses (Figure 2), subsequently lowering the demand for charged tRNAs as it has been observed previously for Leu starvation ^39^.

While aminoacyl-tRNA synthetases (ARSs) are known to occasionally misactivate tRNAs with structurally similar amino acids, some ARSs developed an intrinsic editing activity that ensures extreme specificity under normal conditions ^56^. However, under conditions of BCAA starvation, cells employ adaptive mechanisms to maintain protein synthesis and attempt to reduce ribosome stalling. One such strategy involves adjusting translational fidelity by allowing amino acid misincorporation. While traditionally considered detrimental due to its association with protein malfunction and diseases, emerging evidence suggests that amino acid misincorporation can serve as a conserved adaptive response to environmental stresses, supporting cell survival ^57,58^. Early studies in *E. coli* demonstrated that phenylalanine scarcity led to leucine misincorporation at phenylalanine codons ^59^. A striking example of this adaptive mechanism has also been observed in human cells during Ile starvation. Under such conditions, isoleucyl-tRNA synthetase (IARS1) can misacylate tRNA^Ile^ with valine, resulting in valine incorporation at isoleucine codons ^60^. This substitution helps sustain translation and cellular homeostasis, preventing widespread translational arrest that would otherwise occur if ribosomes stalled at every isoleucine codon. Beyond misincorporation, amino acid starvation can induce also ribosomal frameshifting, leading to the production of aberrant proteins with potential implications for cellular functions ^30,61–63^. Additionally, ribosomes may slide over codons and resume translation downstream ^64^. While not directly related to elongation arrest, it′s worth noting these complex behaviors ribosomes can exhibit under starvation conditions, which might contribute to the observation that some BCAA starvations show much milder elongation changes than others.

Moreover, posttranscriptional modifications of tRNA and mRNA play a crucial role in maintaining the accuracy and efficiency of protein synthesis. In our proteomic analysis, we observed differential expression of OSGEP and MOCS3, enzymes involved in tRNA modifications (Supplementary Data 3). OSGEP is associated with the t^6^A modification at position 37 of ANN-accepting tRNA, which was suggested to be critical for maintaining the correct reading frame during translation ^65–67^. MOCS3 is linked to the mcm^5^s^2^U modification at the wobble position (U34) of tRNA, which enhances codon-anticodon pairing accuracy ^68,69^. In general, modifications in the tRNA anticodon loop have been shown to improve codon recognition and decoding accuracy. Defects in these modifications can increase translational errors and activate stress response pathways, such as the GCN4-dependent expression of general amino acid control (GAAC) genes, even in the absence of amino acid starvation ^70^. Similarly, modifications on mRNA, including N6-methyladenosine (m^6^A), influence mRNA stability, splicing, translation efficiency, and localization ^71^. m^6^A has been shown to be dynamically regulated under stress and to regulate the ATF4 expression and autophagy induction under amino acid starvation ^72,73^. Moreover, m^6^A deposition influences the translation elongation by altering the decoding kinetics by tRNAs and results in occurrence of ribosome stalling ^74,75^. Interestingly, the PCA of our RNA-seq reveals an enrichment of transcripts involved in m^6^A pathway among the transcripts driving the separation of starvations involving ile from the rest. Thus, considering these modifications will help to fully understand the diverse landscape of translation elongation changes under stress, as they directly impact translational fidelity and cellular responses.

Beyond these tRNA dynamics, our data also highlight the importance of the codon positional context within mRNAs, indicating that where a codon is located within the CDS can influence both the extent of ribosomal stalling and overall translation efficiency during nutrient stress. It is well established that codon usage along mRNAs is not random but follows distinct positional patterns. Early work suggests that the 5′ CDS region of highly expressed genes are often enriched in “suboptimal” or rare codons ^76,77^. These regions are thought to serve as regulatory checkpoints to prevent ribosome traffic jams, and allow proper co-translational folding of the nascent peptides ^76,78^, ultimately allowing efficient protein synthesis. In addition, the interplay between codon bias and mRNA secondary structure (modulated by the underlying codon sequence) at the start of transcripts adds another layer to translation regulation ^79,80^. Expanding on these insights, our findings demonstrate that under BCAA limiting conditions, valine codon enrichment at the 5′ end of transcripts can lead to increased ribosome density and early ribosome stalling events. These stalling events may create a bottleneck, e.g. when terminal stalling occurs, that prevents ribosomes from reaching downstream regions of the mRNA, masking other potential stalling sites further along the transcript (i.e. on Ile codons). Alternatively, even if ribosomes do not stall terminally, they might transiently slow down at these early valine codons and reduce the local, immediate demand for amino acids like isoleucine. Our finding that translation inhibition rapidly restores charged tRNA pools across all starvations (Figure 6) supports this model: A local slow down of translation at valine codons may allow aminoacyl-tRNA synthetases additional time to recharge Ile-tRNAs before the ribosome encounters Ile codons downstream, ultimately reduce stalling at downstream Ile codons during triple starvation.

Although we identified ribosomal stalling sites on transcripts with simultaneous decreases in protein synthesis as a key feature of valine deprivation, our study does not directly measure the downstream consequences of the stalled translation on protein degradation, cellular proteostasis, or mRNA stability. Recent work has shown that persistent ribosome stalling can activate ribosome-associated quality control (RQC) pathways, which facilitate nascent chain degradation and can also trigger targeted decay of the associated mRNA ^81,82^. Moreover, ribotoxic stress responses can be initiated by ribosome collisions or prolonged stalling, leading to activation of stress kinases such as JNK and influencing broader proteostatic programs ^83,84^. These mechanisms underscore how terminal stalling or slowed elongation can have far-reaching consequences for both the protein and RNA components of the translational apparatus. However, it remains unclear whether the observed stalling in our BCAA starvations induces terminal stalling, or whether a deceleration of elongation without termination, may cause a differential engagement of RQC and ribotoxic stress pathways. Future studies employing disome (collided-ribosome) sequencing approaches and specialized reporter assays, combined with global proteostasis analyses (e.g., ubiquitin-profiling or nascent chain tracking) and mRNA stability measurements, will be necessary to disentangle the precise nature of ribosome slowdown and its direct impact on protein turnover and quality control mechanisms.

In summary, our integrated analysis reveals a significant codon-specific impact of ribosome stalling on translation during BCAA starvation. These effects are critically modulated by the positional distribution of codons along the mRNA, creating a novel bottleneck phenomenon that underscores an underappreciated regulatory mechanism in cellular adaptation to nutrient stress.

## Data availability

Raw sequencing data can be accessed through the GEO repository, accession numbers: GSE291652 and GSE291653.

## Supplementary Data

Supplementary Figures

Supplementary Material and Methods: Reagents and Oligo sequences

Supplementary Data 1: Dwell time changes upon BCAA starvation

Supplementary Data 2: Free intracellular amino acid levels upon BCAA starvation

Supplementary Data 3: GO terms and DGE analysis of RNA-seq & Ribo-seq and RD and proteomic changes upon BCAA starvation

Supplementary Data 4: Transcript specific polarity scores (Distribution of ribosome density from 5′ to 3′)

Supplementary Data 5: Extracted positions and transcripts displaying ribosome stalling (identified by peak calling)

## Author contributions

LW, CG, and FN conceived the study. LW designed, planned and performed the experiments, analyzed the data, and wrote the manuscript. LW and CG performed the computational analysis. CG provided project oversight and supervision. All authors reviewed, edited and approved the final manuscript.

## Acknowledgements

We thank the EPFL Proteomics Core Facility, particularly Diego Chiappe, for performing the TMT-based quantitative proteomics. We also thank the EPFL Gene Expression Platform for conducting the sequencing experiments. Moreover, we thank Héctor Gallart-Ayala and Julijana Ivanisevic from the UNIL Metabolomics Platform for the support with the intracellular amino acid measurements. Finally, we also thank all members of the Naef lab for their valuable input, discussions, and support throughout this project, in particular Nagammal Neelagandan for conducting a pilot experiment.

## Supplementary Figures

**Supplementary Figure 1.**
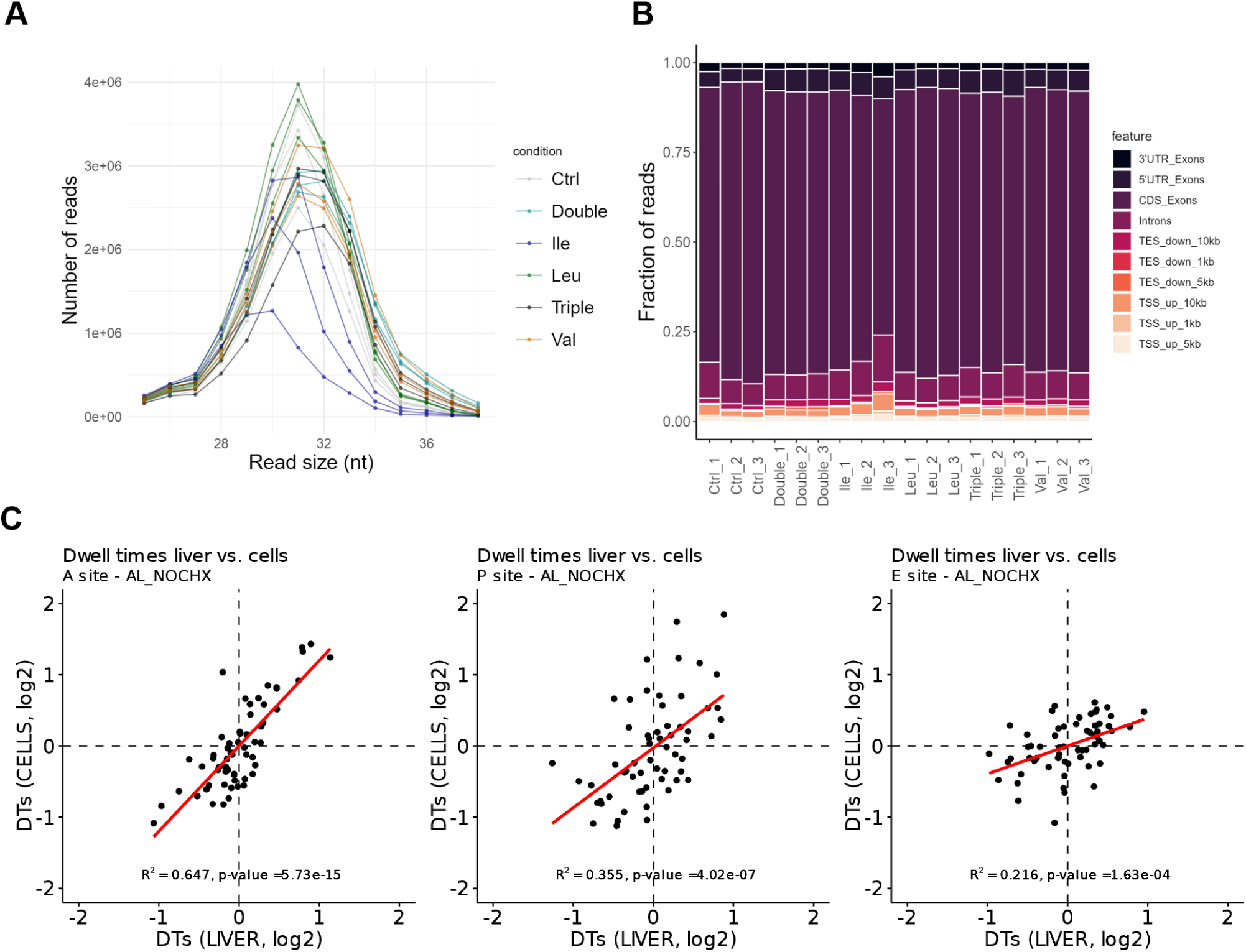
Quality control and comparison of ribosome profiling data across experimental conditions. (A) Footprint length distribution for each sample. (B) Percentage of mapped reads to different transcript features such as CDS and UTRs. (C) Scatter plots comparing dwell times (log₂) in liver ^1^ vs. cells (this study) at the ribosomal A, P, and E sites. Linear regression (red line) with R² and p-value displayed. Dashed lines indicate zero reference points.

**Supplementary Figure 2.**
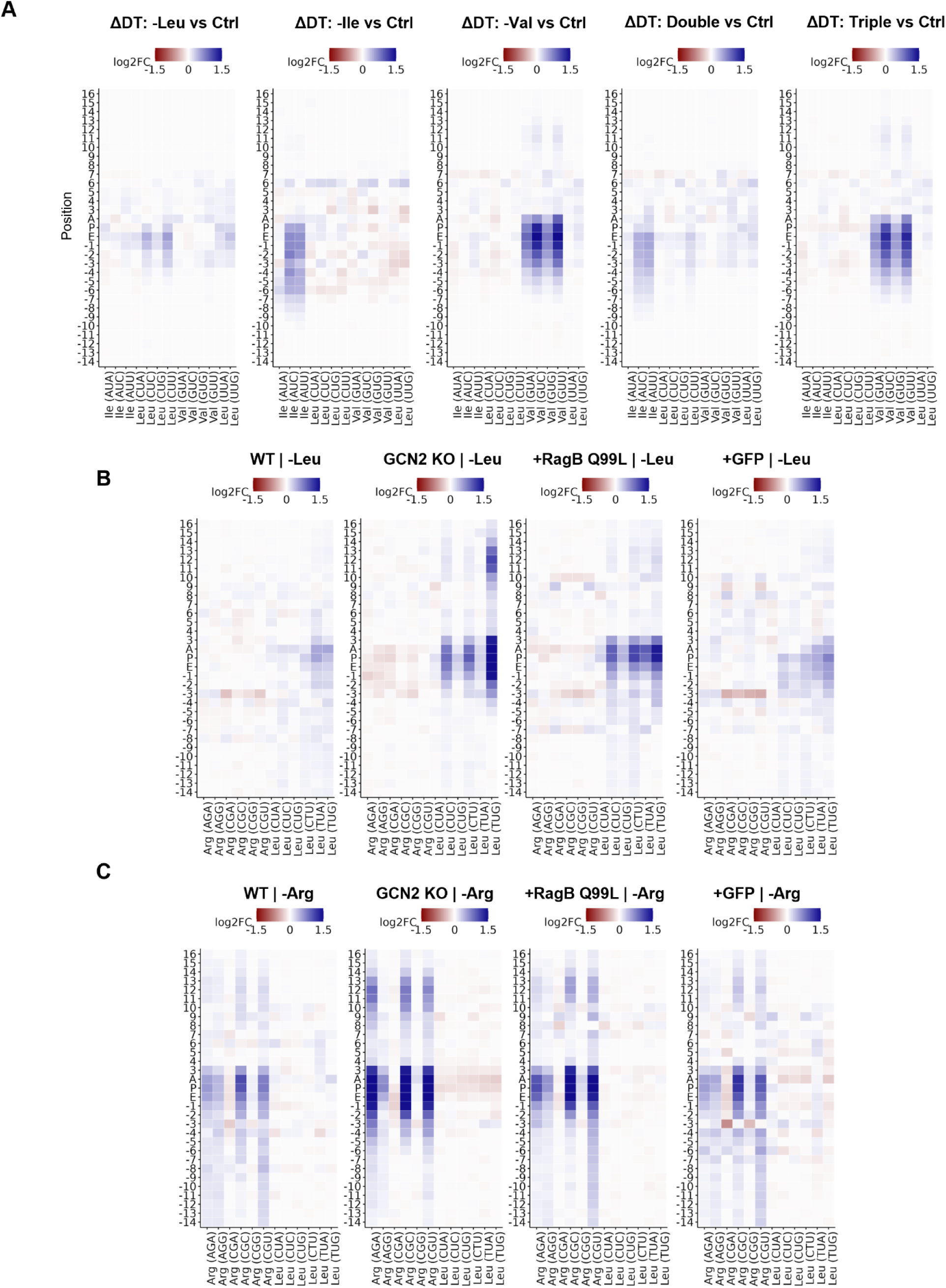
Amino acid starvation alters ribosomal dwell times beyond the A site in mammalian cells. (A-B) Heatmaps of ΔDT (log₂FC) relative to Ctrl for annotated codons across ribosomal positions (−14 to 18, including E, P, and A sites) across (A) BCAA starvations in NIH3T3 cells of this study, (B-C) DTs resulting from reanalysis of published Ribo-seq data ^2^, produced in HEK293T cells (Wildtype, GCNO KO, RagB mutant, GFP overexpression) starved for 6h of (B) leucine or (C) arginine.

**Supplementary Figure 3.**
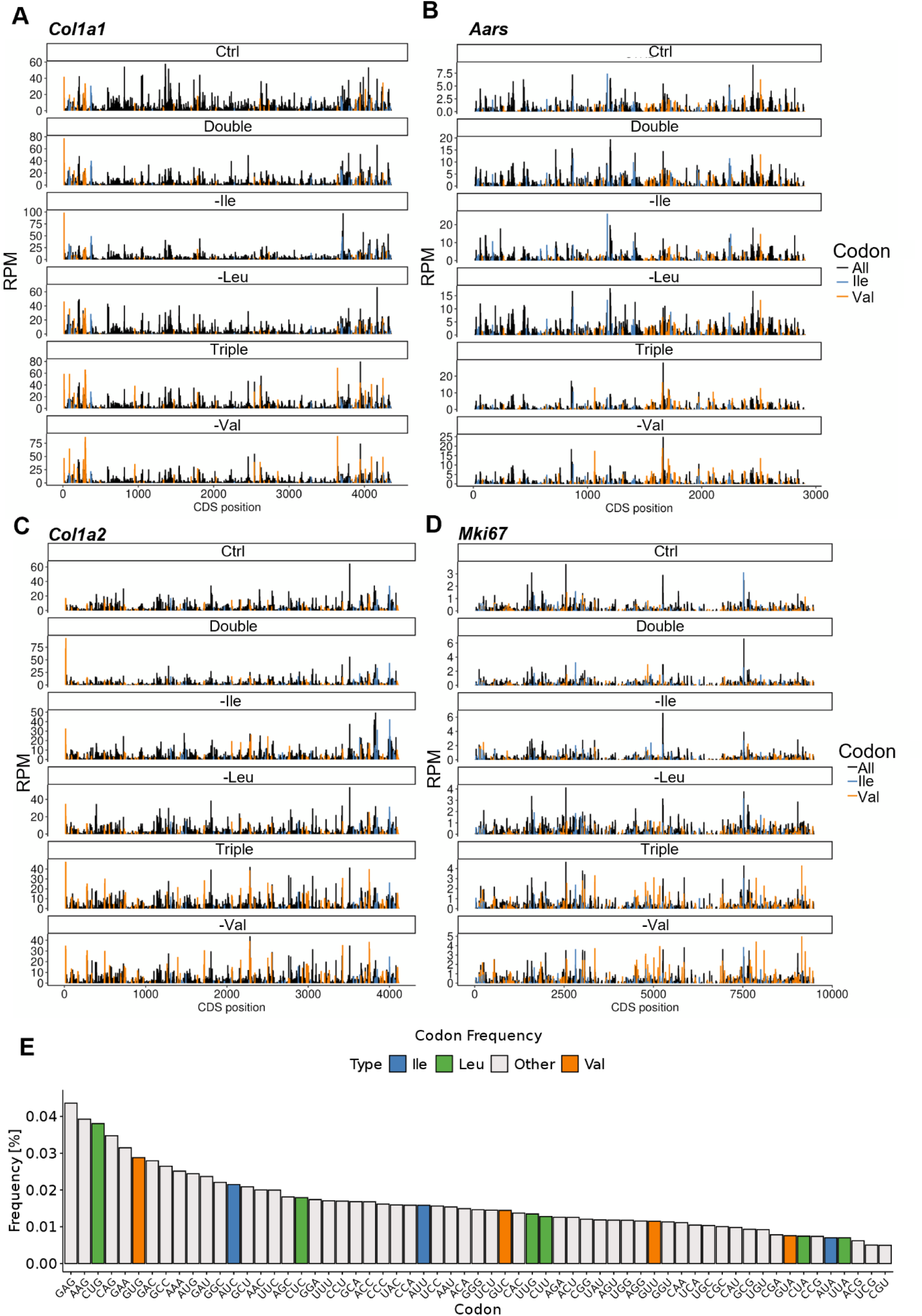
Ribosome density profiles at the gene-level illustrate differential stalling across conditions. **(A-D)** Representative ribosome profiling tracks for the transcripts of (A) *Col1a1*, (B) *Aars*, (C) *Col1a2* and (D) *Mki67* with Val and Ile codon positions (P-site ± 1) annotated. (C) Bar plot showing codon frequencies within transcripts expressed in mouse NIH3T3 cells. Codons for Leu (green), Ile (blue), and Val (orange) are highlighted. Frequencies are averaged across transcripts, and codons are sorted by decreasing frequency.

**Supplementary Figure 4.**
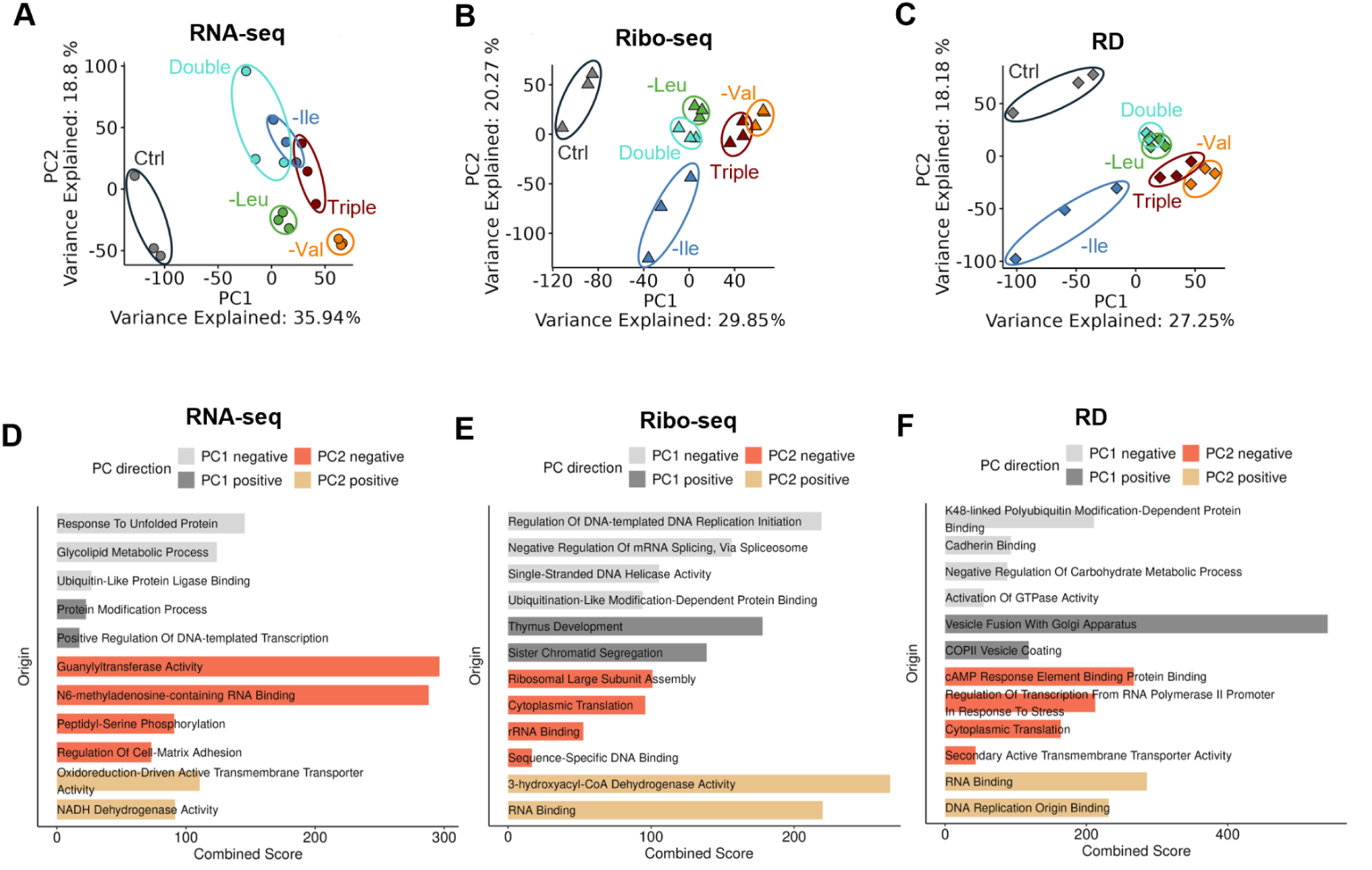
Principal component analysis reveals distinct transcriptional and translational regulation across starvation conditions. (A-C) Principal Component Analysis (PCA) of (A) RNA-seq, (B) Ribo-seq, and (C) ribosome densities (RD) data after group-wise mean-centering relative to Ctrl. PC1 vs. PC2 plot shows sample clustering based on gene expression variation. Colors represent treatment conditions, while shapes indicate data type. Variance explained by each principal component is displayed on the axes. (D-F) Top enriched GO terms for PC1 and PC2 (positive and negative loadings) from PCA analysis of (D) RNA-seq, (E) Ribo-seq and (F) RD. Transcripts included in the enrichment analysis were selected based on loading values exceeding ±15. GO terms were filtered for adjusted p-value < 0.05 and combined enrichment score > 15.

**Supplementary Figure 5.**
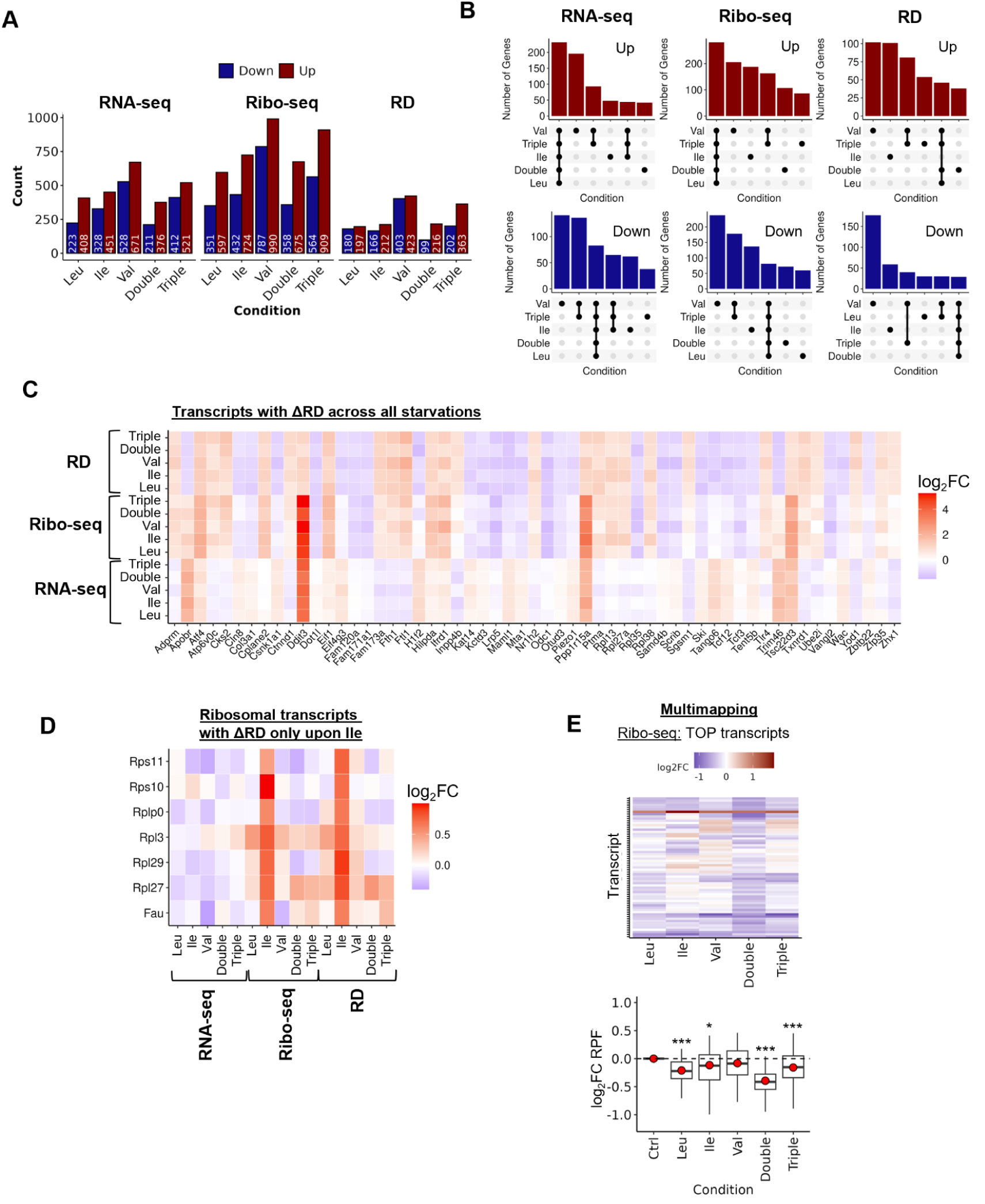
Analysis of transcript-specific ribosome densities (RD) shows condition-specific patterns. (A) Bar plot showing the number of significantly upregulated (red) and downregulated (blue) transcripts across different deprivation conditions (-Leu, -Ile, -Val, Double, Triple). Transcripts were classified as differentially expressed based on |log₂FC| > 0.58 with p < 0.05. Separate counts are shown for RNA-seq, Ribo-seq, and RD data. (B) UpSet plots illustrating the overlap of differentially expressed genes across deprivation conditions for RNA-seq, Ribo-seq, and RD datasets. Genes were considered significantly upregulated, downregulated based on |log₂FC| > 0.58 with p < 0.05. Bars indicate the number of genes shared across multiple conditions. (C) Heatmap displaying log_2_FC of transcripts with dysregulated ribosome density (RD) in all starvation conditions (upregulated or downregulated). (D) Heatmap displaying log_2_FC of a subset of transcripts that only show a significant RD change upon Ile starvation. Displayed are the transcript related to ribosomal proteins. (E) Heatmap and quantification of log₂FC for the known TOP motif containing transcripts^3^ in our Ribo-seq using unique and multimapping reads (see Material and Methods). Genes are clustered based on their expression profiles using k-means clustering (k = 12). In the boxplot, red points indicating the mean expression change. Significance was assessed using unpaired t-tests.

**Supplementary Figure 6.**
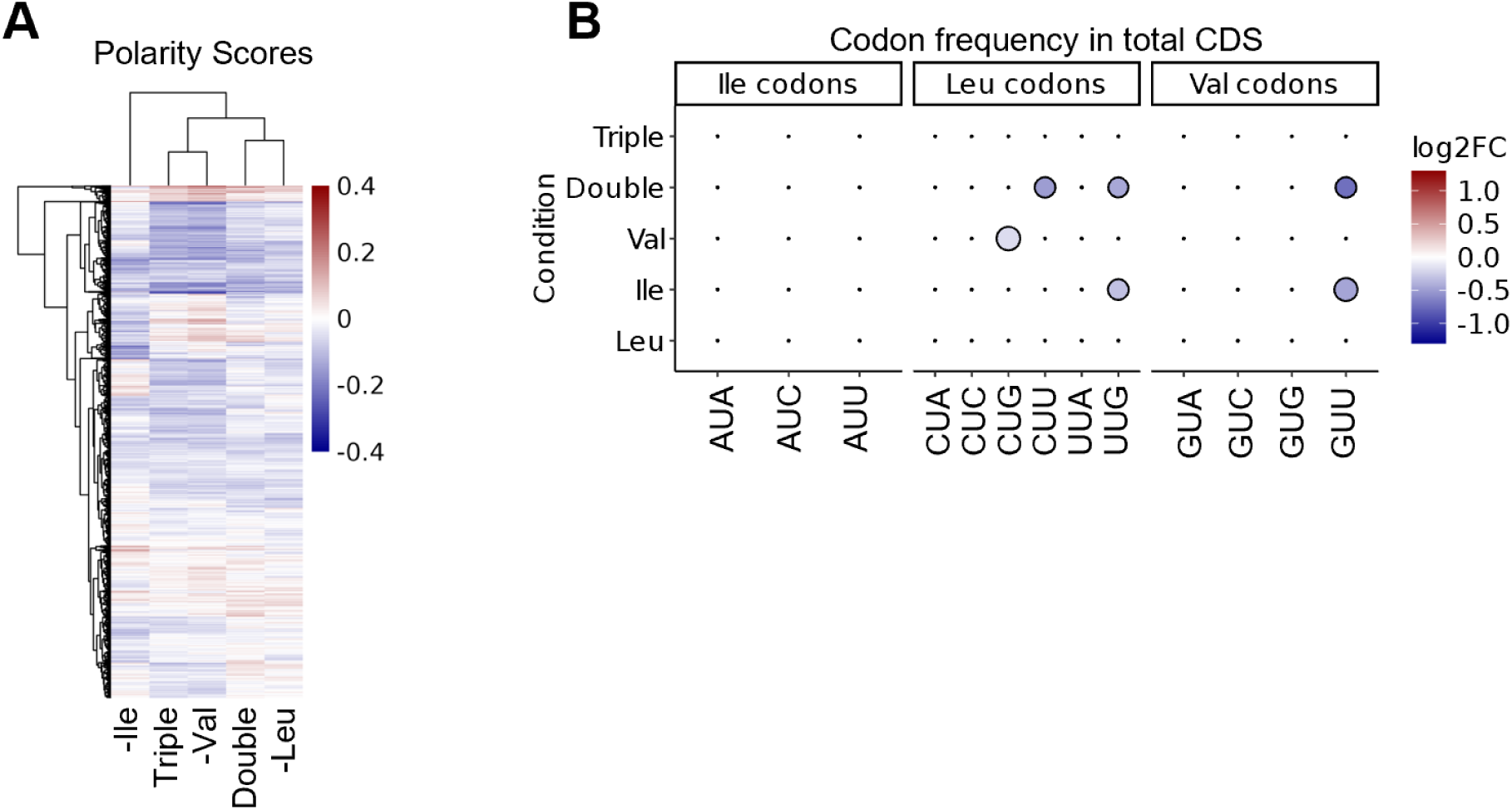
Transcripts with polarity score shifts do not have an increased frequency of starved amino acid in total CDS. (A) Heatmap of transcript-specific polarity score changes across the tested starvations relative to Ctrl. (B) Dot plot summarizing changes in normalized codon frequencies in groups of transcripts extracted in Figure 4D in their total CDS. For each codon group (Val, Leu, and Ile codons), the log_2_FC relative to all transcripts is plotted. Dot size reflects the -log₁₀(p-value) from unpaired t-tests (with BH adjustment), and the fill color indicates the magnitude and direction of change.

**Supplementary Figure 7.**
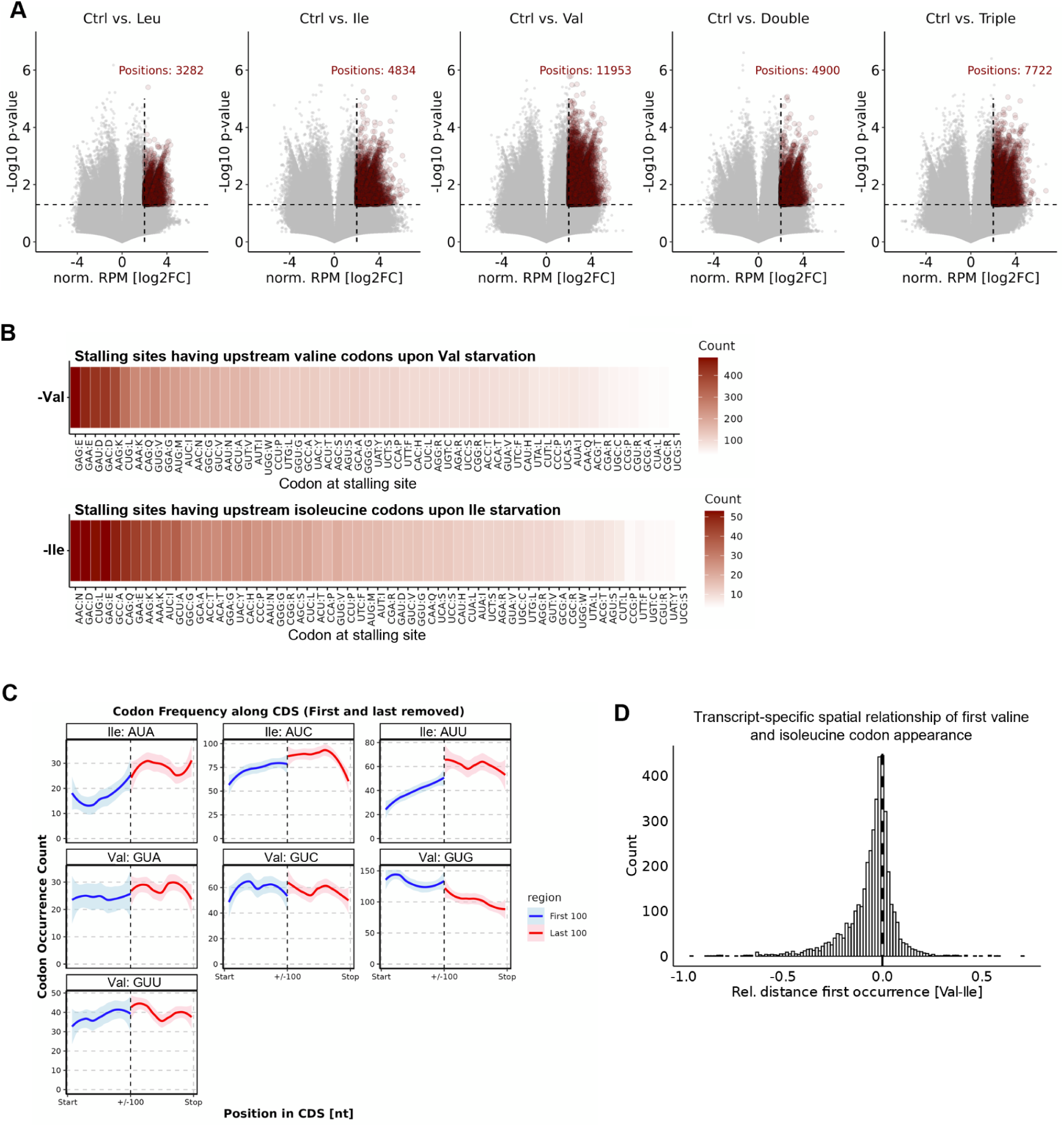
Non-uniform codon distribution creates elongation bottlenecks. (A) Volcano plots displaying log₂FC versus -log₁₀(p-value) for each position within transcripts. Positions meeting peak-calling criteria for significant upregulation are highlighted in red. (B) Heatmap illustrates the frequency of specific codons at stalling sites, conditioned on the presence of valine (Val) or isoleucine (Ile) codons in the 1–3 positions upstream of the stalling site. (C) Frequencies of isoleucine (Ile) and valine (Val) codons in the first and last 100 codons across all transcripts. (D) Spatial analysis of occurrence of first Val codon relative to first Ile codon on each transcript.

**Supplementary Figure 8.**
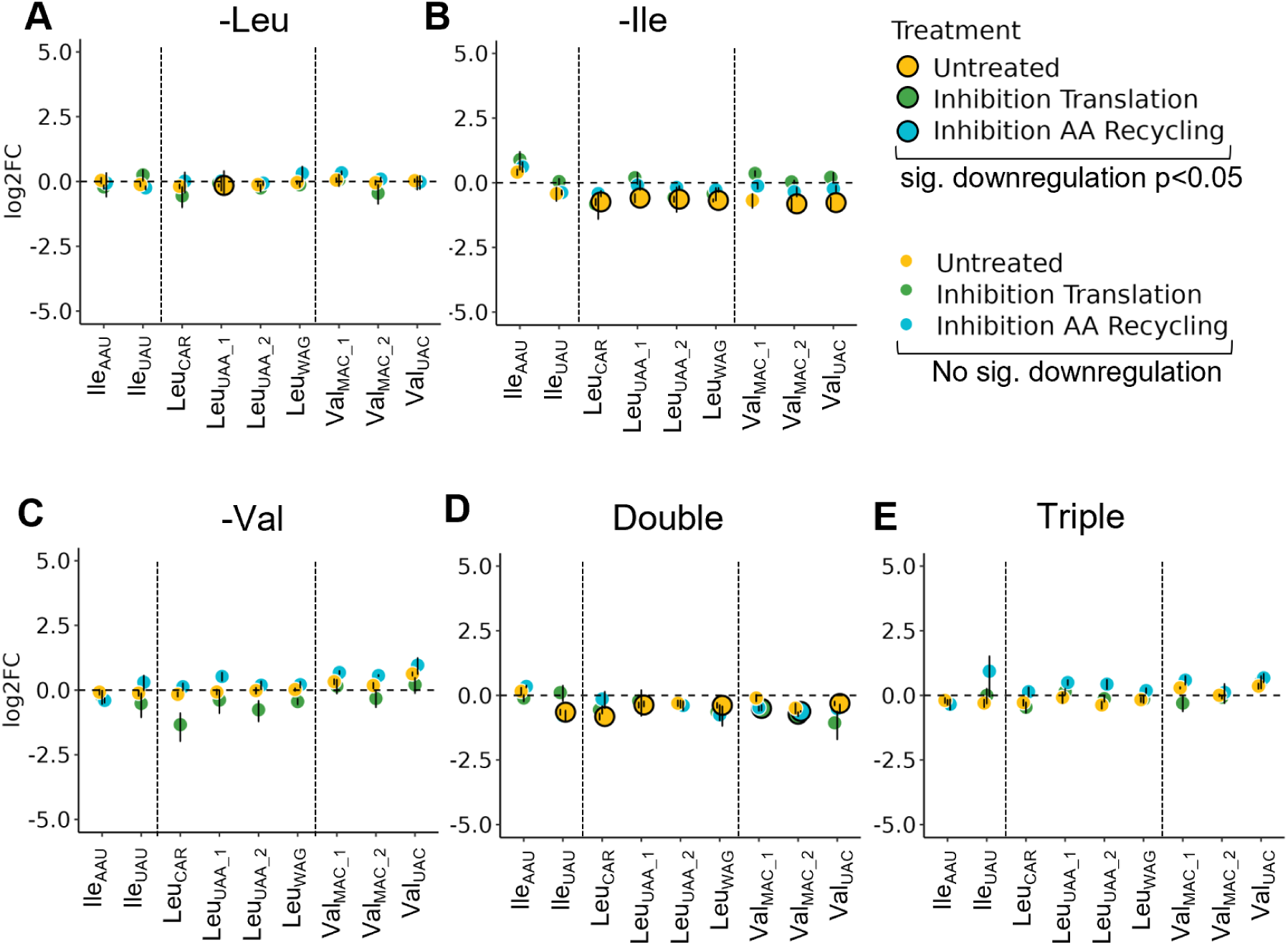
Total tRNA levels show minimal changes across starvation conditions. (A–E) Differential tRNA isoacceptor level under the indicated starvation conditions. Yellow indicates starved cells without treatment. Green represents cells treated with 100 µg/mL cycloheximide (CHX) in the last 30 min of starvation to inhibit translation. Blue represents cells treated with 10 µM MG132 and 160 nM Bafilomycin A1 to inhibit proteasomal and autophagic degradation (amino acid recycling inhibition). Certain tRNA isodecoders could not be measured separately and are therefore represented with IUPAC ambiguity codes: W (A or U), R (A or G), and M (A or C). Large outlined dots indicate significant downregulation. Sample sizes: untreated (n = 5), translation and AA recycling inhibition (n = 2).

